# Tumor- and Nerve-Derived Axon Guidance Molecule Promotes Pancreatic Ductal Adenocarcinoma Progression and Metastasis through Macrophage Reprogramming

**DOI:** 10.1101/2023.10.24.563862

**Authors:** Noelle R. J. Thielman, Vanessa Funes, Sanjana Davuluri, Hector E. Ibanez, Wei-Chih Sun, Juan Fu, Keyu Li, Stephen Muth, Xingyi Pan, Kenji Fujiwara, Dwayne Thomas, MacKenzie Henderson, Selina Shiqing Teh, Qingfeng Zhu, Elizabeth Thompson, Elizabeth M. Jaffee, Alex Kolodkin, Fengxi Meng, Lei Zheng

## Abstract

Axon guidance molecules were found to be the gene family most frequently altered in pancreatic ductal adenocarcinoma (PDA) through mutations and copy number changes. However, the exact molecular mechanism regarding PDA development remained unclear. Using genetically engineered mouse models to examine one of the axon guidance molecules, semaphorin 3D (SEMA3D), we found a dual role for tumor-derived SEMA3D in malignant transformation of pancreatic epithelial cells and a role for nerve-derived SEMA3D in PDA development. This was demonstrated by the pancreatic-specific knockout of the *SEMA3D* gene from the *KRAS^G12D^* and *TP53^R^*^172^*^H^* mutation knock-in, *PDX1-Cre* (KPC) mouse model which demonstrated a delayed tumor initiation and growth comparing to the original KPC mouse model. Our results showed that SEMA3D knockout skews the macrophages in the pancreas away from M2 polarization, providing a potential mechanistic role of tumor-derived SEMA3D in PDA development. The KPC mice with the SEMA3D knockout remained metastasis-free, however, died from primary tumor growth. We then tested the hypothesis that a potential compensation mechanism could result from SEMA3D which is naturally expressed by the intratumoral nerves. Our study further revealed that nerve-derived SEMA3D does not reprogram macrophages directly, but reprograms macrophages indirectly through ARF6 signaling and lactate production in PDA tumor cells. SEMA3D increases tumor-secreted lactate which is sensed by GPCR132 on macrophages and subsequently stimulates pro-tumorigenic M2 polarization in vivo. Tumor intrinsic- and extrinsic-SEMA3D induced ARF6 signaling through its receptor Plexin D1 in a mutant KRAS-dependent manner. Consistently, RNA sequencing database analysis revealed an association of higher *KRAS^MUT^* expression with an increase in *SEMA3D* and *ARF6* expression in human PDAs. Moreover, multiplex immunohistochemistry analysis showed an increased number of M2-polarized macrophages proximal to nerves in human PDA tissue expressing SEMA3D. Thus, this study suggests altered expression of SEMA3D in tumor cells lead to acquisition of cancer-promoting functions and the axon guidance signaling originating from nerves is “hijacked” by tumor cells to support their growth. Other axon guidance and neuronal development molecules may play a similar dual role which is worth further investigation.

**One sentence summary:** Tumor- and nerve-derived SEMA3D promotes tumor progression and metastasis through macrophage reprogramming in the tumor microenvironment.

**STATEMENT OF SIGNIFICANCE:** This study established the dual role of axon guidance molecule, SEMA3D, in the malignant transformation of pancreatic epithelial cells and of nerve-derived SEMA3D in PDA progression and metastasis. It revealed macrophage reprogramming as the mechanism underlying bothroles. Together, this research elucidated how inflammatory responses promote invasive PDA progression and metastasis through an oncogenic process.

## INTRODUCTION

Pancreatic ductal adenocarcinoma (PDA) is a dismal malignant disease, as the third leading cause of cancer-related death with the lowest survival rate compared to any other major cancer type at 12%^1^. This poor prognosis is attributed to already advanced metastatic disease upon diagnosis, precluding surgical intervention, and a high systemic recurrence rate even following surgical resection^2^. The tumor microenvironment (TME) is recognized for its essential role in promoting progression and metastasis in PDA. Nerves in the TME, and associated axon guidance genes they express, have been increasingly appreciated for their role in influencing disease progression^3–7^. Perineural invasion (PNI), the neoplastic invasion of tumor cells into or surrounding nerves, is a characteristic feature in PDA, occurring in 80-100% of human PDA, and is associated with poor patient prognosis and aggressive tumor characteristics^8–10^. We previously observed that the nerve-tumor cell signaling involved in PNI is linked to increased metastasis^10^.

The expression of axon guidance molecules has been found to be most frequently altered in PDA through mutations and copy number changes^11^. In a recent single-nucleus and spatial transcriptome profiling study of PDA, a panel of neuronal development genes including axon guidance molecules were found to be associated with PNI and poor prognosis^12^. These proteins comprise several families, including semaphorins and their plexin receptors, which have been found to exhibit elevated expression in the TME of several cancer types, including PDA, and to promote disease progression^10,13–16^. Axon guidance proteins have also been found to play a critical role in communication between several cell types in the PDA TME through induction of signaling cascades between epithelial cells, neurons, fibroblasts, and macrophages^17–20^. Our previous analysis of signaling pathways regulated by Annexin A2, a PDA-associated tumor antigen and metastasis-related protein, identifies the secreted protein Semaphorin 3D (SEMA3D) and its co-receptors, Plexin D1 (PLXND1) and neuropilin-1, as being involved in increasing nerve migration and PDA cell invasiveness through both autocrine and paracrine signaling to promote PDA progression and metastasis^10,15,21,22^. However, the precise molecular mechanisms underlying this signaling event remain unknown, particularly regarding whether altered axon guidance molecule expression in tumor cells promotes cancer or axon guidance signaling from intratumoral nerves is “hijacked” by tumor cells to support tumor cell growth and metastasis.

Axon guidance molecules, including semaphorins, also play important roles in immune cell regulation and can, therefore, influence the immune landscape in PDA. Semaphorins promote lymphocyte proliferation, inflammatory responses, and immune cell chemotaxis^23–25^. Macrophages are also a dynamic and influential cell type in the progression or suppression of PDA. Classically activated M1-like macrophages produce proinflammatory cytokines and inhibit tumor cell proliferation while alternatively activated M2-like macrophages are considered anti- inflammatory and promote tumor cell proliferation^26,27^. TME signaling can reprogram macrophages to become M2-like macrophages, which can both support the progression and metastasis of PDA^28,29^ and influence the efficacy of immunotherapy treatments^30,31^. However, the impact of tumor- or nerve-derived SEMA3D on tumor progression and metastasis through immune cell reprogramming is not known. Therefore, using a genetically engineered mouse model of PDA, we tested the hypothesis that both tumor- and nerve-derived SEMA3D promote tumor progression and metastasis through macrophage reprogramming.

## RESULTS

### KPCS mouse model demonstrates a delayed tumor initiation and growth compared to KPC mouse model

We previously identified a role for tumor-derived SEMA3D in perineural invasion^10^. To assess the role of SEMA3D in the development, progression, and metastasis of PDA, a KPC-SEMA3D knock out mouse model was created. The KPC mouse model is a genetically engineered mouse model of PDA, previously established through a pancreatic-specific knock-in of conditional alleles of the *KRAS^G12D^* and *TP53^R^*^172^*^H^* mutations driven by Cre recombinase with a *PDX1* promotor and backcrossed to a C57Bl/6 background^32^. Mice with conditional expression of SEMA3D, in which the *SEMA3D* gene is flanked by loxP sites, were crossed with KPC mice. Therefore, mutations causing a loss-of-function (LOF) of SEMA3D and TP53, and a KRAS gain-of-function (GOF) are specifically activated in the pancreas under the PDX-1 promoter, generating *KRAS^G12D^ TRP53^R^*^172^*^H^ SEMA3D^fl/fl^ PDX-1-Cre* mice (designated KPCS mice). Loss of SEMA3D expression in the pancreatic epithelia was confirmed via immunohistochemical staining of PDA samples from KPCS mice (Figure 1a).

**Figure 1:**
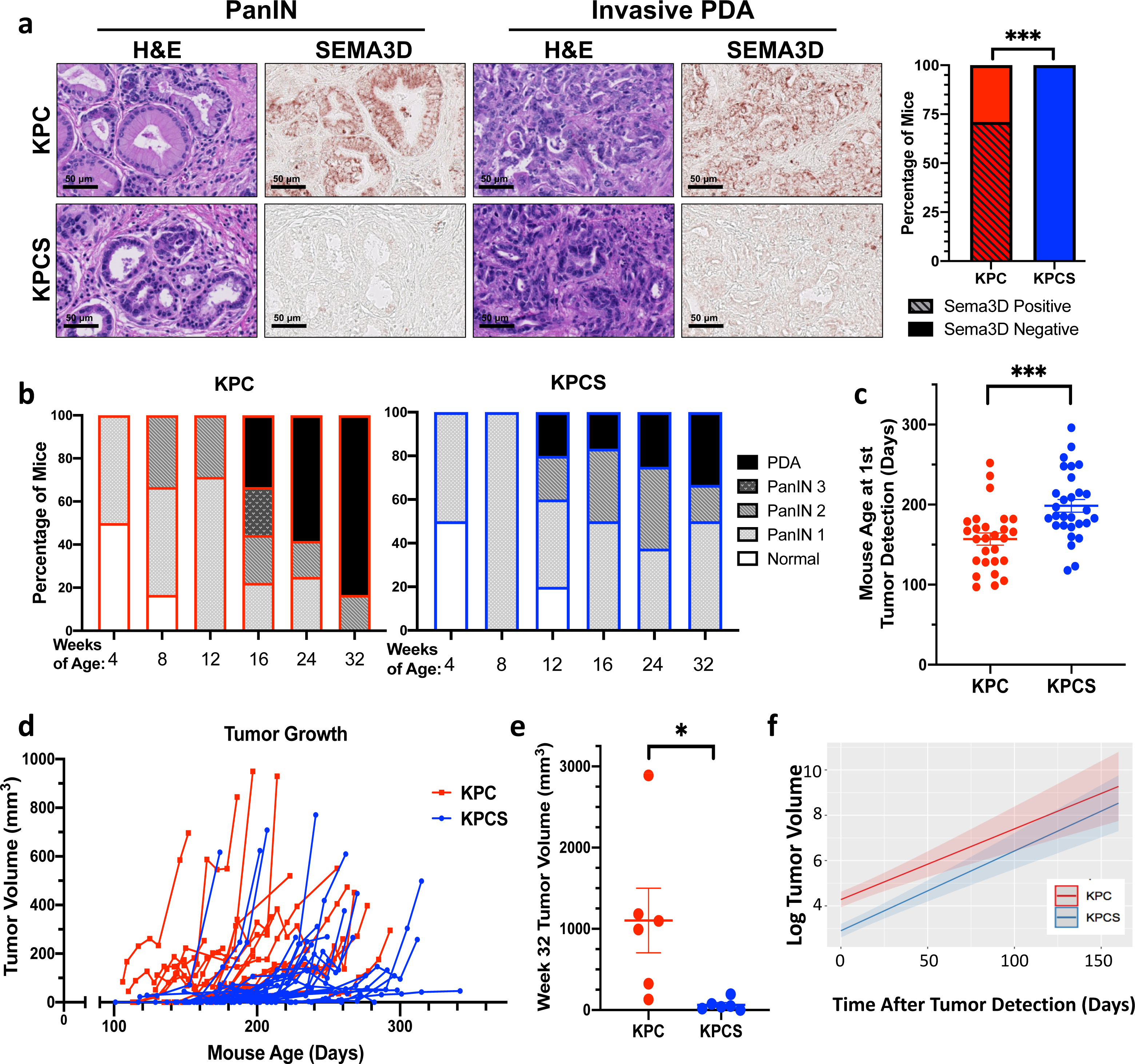
KPCS mouse model demonstrates a delayed tumor initiation and growth compared to KPC mouse model. a. Representative SEMA3D and hematoxylin and eosin (H&E) immunohistochemical staining of PanIN and invasive PDA samples from KPC (n=17) and KPCS (n=11) mice along with percentage of mice expressing SEMA3D-positive tumor cells. b. Percentage of KPC and KPCS mice with normal, PanIN1, PanIN2, PanIN3 or PDA at various ages. c. Tumor growth curves starting from the time of initial tumor detection in KPC and KPCS mice. d. Spaghetti plots of tumor volumes measured by the small animal ultrasound for KPC (n = 19) and KPCS (n= 29) mice. e. Tumor size of KPC and KPCS mice at age of 32 weeks. f. Predicted tumor growth curves for KPC and KPCS mice using log transformed tumor volumes measured by ultrasound.

The KPC mouse model develops pancreatic intraepithelial neoplasia (PanIN) lesions that progress stepwise to invasive PDA development analogous to human disease^32^. Similarly, histological analysis by pathologist (E.M.T.) confirmed the stepwise disease progression of PDA in the KPCS mice; however, KPCS mice demonstrated a delayed tumor initiation and development compared to KPC mice of similar ages. The percentage of mice with PDA, compared to other stages of disease, is statistically higher in KPC mice compared to KPCS mice at ages of 16, 24, and 32 weeks combined, by (Fisher’s Exact Test, p<0.05; Figure 1b). Furthermore, we used small animal ultrasound technologies to detect and measure tumor growth throughout the lifetime of KPC and KPCS mice starting at age of 12-16 weeks. KPCS mice were significantly older upon initial tumor detection compared to KPC mice (Figure 1c). In addition, spaghetti plots of individual mouse tumor growth over time demonstrate that KPCS mice have a delayed primary tumor development compared to KPC mice (Figure 1d). Furthermore, primary PDA samples isolated from mice at age of 32 weeks were significantly smaller in KPCS mice compared to KPC mice (Figure 1e).

To examine the tumor growth rate in KPC and KPCS mice, the tumor volume and log tumor volume of each mouse were analyzed starting on the initial day of tumor detection (Figure S1a and S1b). A linear mixed-effects model analysis with log-transformed tumor volume as the dependent variable led to the conclusion that there is a significantly smaller tumor volume in KPCS mice, particularly during the first 50 days, compared to KPC mice, but no difference in overall tumor growth rate (Figure 1f). This result further suggests that SEMA3D plays a role in controlling PDA growth and development; however, a compensatory mechanism arises in KPCS mice to allow PDA tumor growth eventually.

### KPCS mouse model exhibits an increased survival and an absence of metastasis compared to KPC mouse model

To determine the role of SEMA3D in the survival of KPC mice, we monitored 20 KPC and 29 KPCS mice until time of death (Figure 2a). A Mantel-Cox log rank survival analysis was performed and KPCS mice were found to have a statistically significant increase in overall survival, with a median survival of 264 days compared to 213 days in KPC mice. The Mantel-Haenszel hazard ratio was 2.068 when comparing KPC mice to KPCS mice. At time of death, the lung, liver, diaphragm, and intestines were collected to examine extent of metastatic spread in KPC and KPCS mice. All mice were confirmed to have invasive PDA; however, metastatic tumor presence was histologically identified in 76.5% of KPC mice and 0% of KPCS mice (Figure 2b). Furthermore, we used a preclinical model of hepatic metastasis via hemispleen tumor cell injection to examine differences in metastasis formation, as previously described^15,22^. Isogenic tumor cell lines created from KPC and KPCS primary tumor samples were injected into wild-type (WT) mice. Mice receiving KPCS cells via hemispleen injection exhibited statistically increased survival compared to mice receiving KPC cells, further supporting a role for SEMA3D in PDA metastasis (Figure 2c).

**Figure 2:**
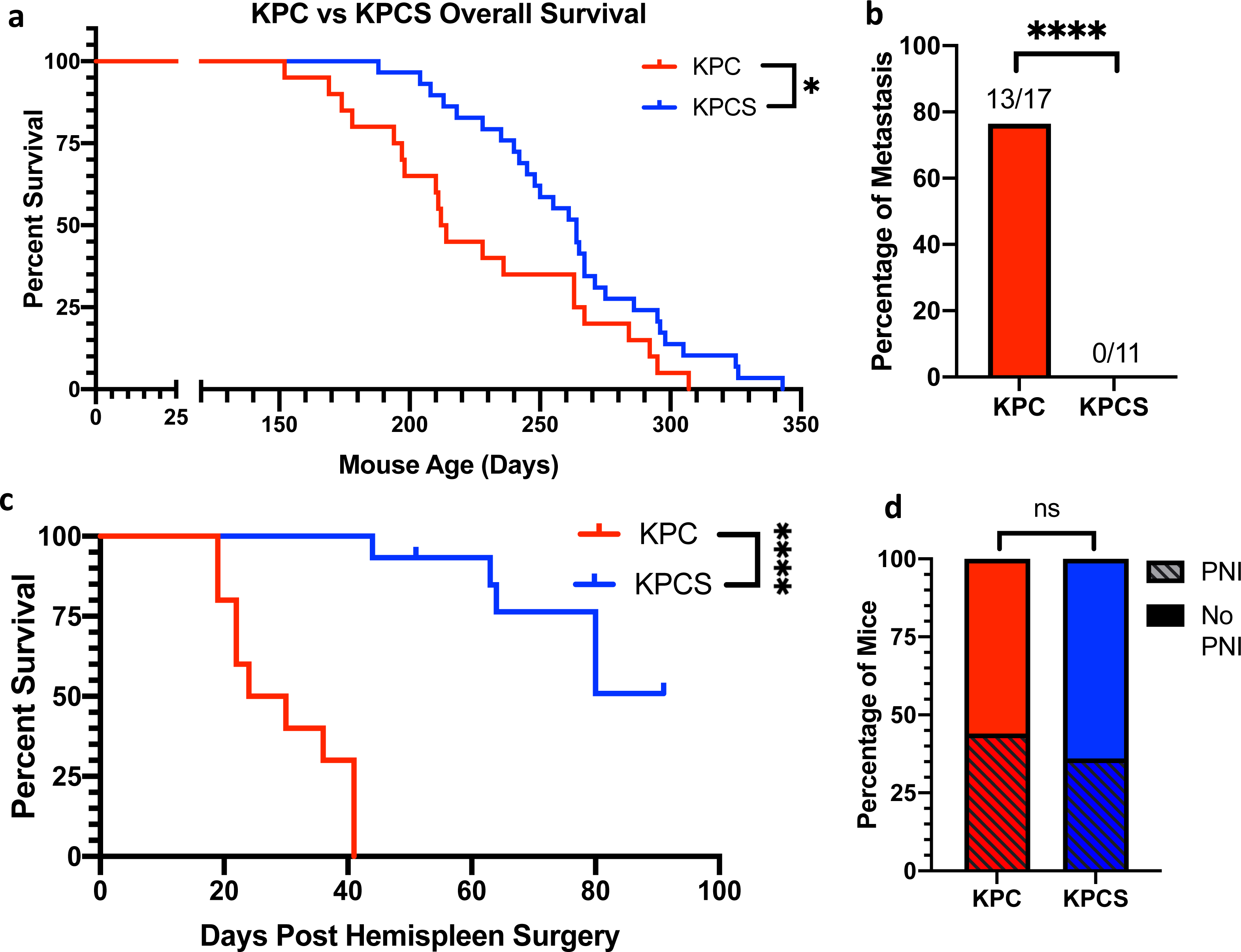
KPCS mouse model exhibits increased overall survival and an absence of metastasis compared to KPC mouse model. a. Overall survival of KPC (n=20) and KPCS (n=29) mice were monitored from time of birth. b. Percentages of KPC (n=17) and KPCS (n=11) mice with metastasis identified in the lung, liver, diaphragm, and/or intestines. c. Survival curve of mice subjected to KPC (n=10) or KPCS (n=15) cells through Hemispleen injection. d. Percentages of KPC (n=18) and KPCS (n=11) mice with histologically confirmed PNI.

Previously, we demonstrated that perineural invasion (PNI), the invasion of tumor cells into nerve sheaths, is a potential SEMA3D-mediated mechanism of metastatic spread in pancreatic cancer^10^. Therefore, H&E stained slides of KPC and KPCS primary tumors at the survival endpoint were analyzed by a pathologist to identify the presence of PNI. KPCS tumors did have slightly less PNI overall, averaging 36% of mice with PNI compared to 44% in KPC tumors (Figure 2d). TUJ1 immunohistochemistry was used to identify intratumoral nerves (Figure S2a). Nerve densities were calculated according to the number of TUJ1-marked nerve bundles within the tumor area, revealing a trend towards increased nerve density in KPC tumors compared to KPCS tumors (Figure S2b). In contrast, we previously found that knockdown of SEMA3D from the KPC tumor cell line significantly reduced nerve density in the orthotopic tumors formed in this model cell line^10^. Here, we found that KPCS mice, compared to KPC mice at the survival endpoint, exhibited only a modest reduction in PNI and intratumoral nerve density, suggesting that other axon guidance pathways or PNI mechanisms may serve as a compensatory fashion to drive the metastatic process if mice did not die from primary tumor growth. Nevertheless, there must be another compensation mechanism to promote this primary tumor growth.

### SEMA3D does not reprogram macrophages directly

We thus hypothesized that such a potential compensation mechanism could result from SEMA3D expression in the stroma compartment. SEMA3D expression in stroma cells was observed to be more prominent in the KPCS tumors as compared to KPC tumors (Figure S3a). Since macrophage reprogramming has been reported to play a role in promoting metastasis and PLXND1, a SEMA3D receptor, may be expressed by macrophages, we next investigated whether SEMA3D has an impact on macrophage reprogramming in the stroma. Bone marrow derived macrophages (BMDMs) were first examined for PLXND1 protein expression. BMDMs were grown and maintained unpolarized (M0) or polarized with M1 or M2 inducing cytokines and stimulants. No PLXND1 protein was detected in M0, M1, or M2 polarized macrophages by western blot, suggesting that BMDMs do not express PLXND1 and SEMA3D does not directly bind to BMDMs (Figure S3b). Furthermore, there was no consistent increase in M1 or M2 gene expression markers in unpolarized M0 (Figure S3a and S3b) or in polarized M1 and M2 cells treated with recombinant SEMA3D protein (SEMA3D-AP) as compared to control recombinant protein (CTRL- AP) (Figure S3c, and S3d). These results suggest SEMA3D signaling in the stroma from secreted SEMA3D produced by tumor cells or stromal cells is unlikely to reprogram macrophages directly, although we cannot rule out the possibility that SEMA3D can reprogram macrophages through other receptors or that PLXND1 may be expressed in tissue-resident macrophages.

### SEMA3D reprograms macrophages indirectly through ARF6 signaling and lactate production in PDA tumor cells

Next, we investigated the possibility that stroma-derived SEMA3D can reprogram macrophages through tumor cells. Previously, we showed that tumor cells can reprogram macrophages through an epigenetic and metabolic reprogramming process^33^. We first confirmed that KPC tumor cells expressed PLXND1 (Figure S3b). Next, we investigated whether or not SEMA3D is able to activate plexin receptor signaling pathways, including those utilizing ARF6, AKT and p38 signaling^34–36^. Recombinant SEMA3D treatment did not cause increased phosphorylation of AKT or p38 (Figure S4a). However, recombinant SEMA3D treatment of KPC cells did result in an increase in levels of the active form of ARF6 protein, ARF6-GTP (Figure 3a) as compared to control-treated cells.

**Figure 3:**
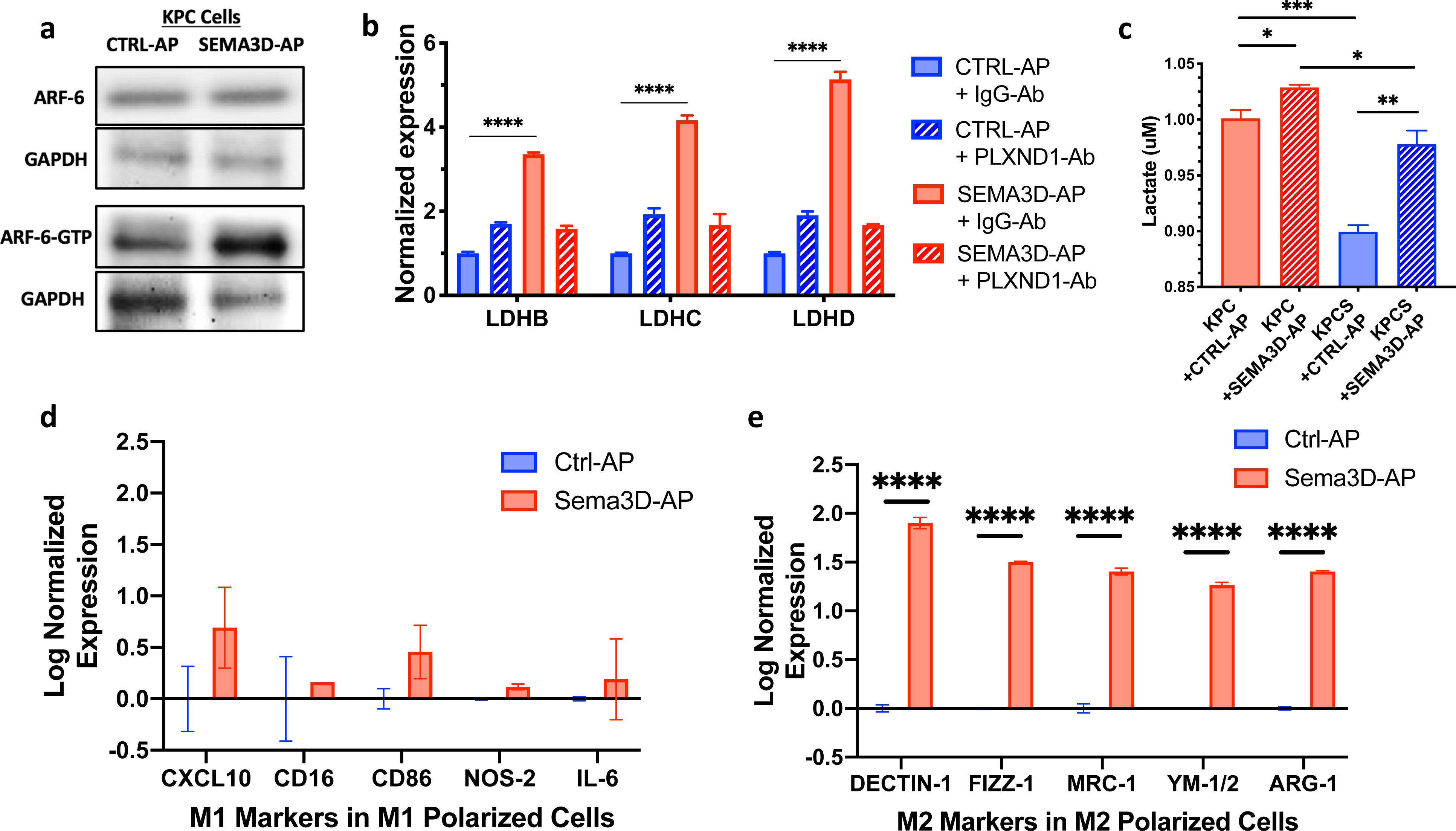
SEMA3D reprograms macrophages indirectly though ARF6 signaling and lactate production in PDA tumor cells a. Western blot analysis of the expression of ARF6 and active ARF6-GTP protein in CTRL-AP and SEMA3D-AP treated KPC tumor cells. b. qRT-PCR analysis of the expression of *LDHB*, *LDHC*, and *LDHD* genes in CTRL-AP or SEMA3D-AP treated KPC cells which were first pre-treated with anti- PLXND1 neutralizing antibody or isotype IgG control antibody. c. Lactate concentration in supernatant from KPC and KPCS cells treated with CTRL-AP or SEMA3D-AP, respectively. d,e. qRT-PCR analysis of gene expression markers for M1 (d) and M2 (e) polarization, respectively, in BMDM cells polarized with M1 or M2 polarizing cytokines and subsequently co-cultured with CTRL-AP or SEMA3D-AP treated KPC cells in the double-chamber.

ARF6 signaling is involved in integrin signaling and intracellular trafficking in angiogenesis through the action of SEMA3E^34,37^, an additional PLXND1 ligand. Recently, ARF6 signaling has also been found to regulate the metabolism of PDA through the Warburg effect; ARF6 stimulated by mutant KRAS is involved in increasing the glycolytic pathway in human PDA cell lines^38^. Therefore, we assessed PLXND1-dependency for SEMA3D-induced ARF6 activation on glycolysis-mediated gene expression in KPC tumor cells. KPC cells were pretreated with a neutralizing antibody (Ab) directed against PLXND1, or IgG Ab, followed by the addition of SEMA3D-AP or CTRL-AP media KPC tumor cells in culture. Quantitative RT-PCR demonstrated a statistically significant increase in the expression of lactate dehydrogenase genes B, C, and D after SEMA3D-AP treatment, and this was blocked by PLXND1 neutralizing antibody pretreatment (Figure 3b), suggesting a PLXND1-dependent increase in glycolytic gene expression following SEMA3D treatment. Furthermore, we collected supernatant from KPC and KPCS cells treated with SEMA3D-AP or CTRL-AP to measure lactate levels after 48 hours of culture, resulting in the observation that KPC cells treated with SEMA3D-AP produced significantly more lactate than KPC cells treated with CTRL-AP protein. CTRL-AP treated KPC cells produced significantly more lactate than CTRL-AP treated KPCS cells; this difference was diminished after the addition of recombinant SEMA3D-AP protein (Figure 3c). Interestingly, SEMA3D-AP treated KPC cells produced significantly more lactate than SEMA3D-AP treated KPCS cells, and this may be due to the decreased levels of PLXND1 expression observed in KPCS cells (Figure S3b). Taken together, these data suggest that exogenous SEMA3D increases glycolytic gene expression and lactate secretion in PDA tumor cells regardless of whether or not tumor cells express endogenous SEMA3D, and this may compensate for the loss of SEMA3D in tumor epithelia in KPCS mice.

Lactate in the TME has previously been shown to modulate macrophage polarization and thus can mediate macrophage reprogramming^33,39^. We examined the changes in characteristic M1 and M2 gene expression markers that occurred in the M0 macrophages after treatment with 25nM lactate containing media. These lactate-treated non-polarized M0 macrophages exhibited significant decreases in M1 expression genes and modest upregulation of M2 expression genes compared to control macrophages (Figure S4b).

Next, we utilized an *in vitro* co-culture model to mimic the TME of PDA by culturing BMDMs with tumor cells treated with exogenous recombinant SEMA3D-AP or CTRL-AP. BMDMs were grown, polarized with M1 or M2 inducing-cytokines and then seeded on the bottom chamber of a transwell co-culturing system. KPC cells treated with CTRL-AP or SEMA3D-AP were then seeded on the top chamber of the transwell system. The co-culturing chamber was separated by one-micron pores, allowing media to be shared between the cell types with no direct contact between polarized BMDMs and KPC cells, which eliminated the possibility of previously described direct-contact mediated reprogramming of macrophages^33^. In these experiments no significant M1 gene expression changes were observed in polarized M1 cells cultured with KPC cells in either CTRL-AP or SEMA3D-AP containing media (Figure 3d). However, M2-polarized cells co-cultured with KPC cells treated with SEMA3D-AP exhibited significantly higher expression of M2 gene markers as compared to M2-polarized cells co-cultured with CTRL-AP treated KPC cells (Figure 3e). In contrast, when M1 or M2 polarized macrophages were co-cultured with KPCS cells, there was no consistent increase in M1 or M2 gene expression markers, likely due to the reduction of PLXND1 protein expression in KPCS cells (Figure S4c). This result suggests that the role of SEMA3D in macrophage reprogramming through PDA tumor cells is PLXND1-dependent even though exogenous SEMA3D is able to induce lactate production in KPCS tumor cells.

### SEMA3D-induced lactate is sensed by GPCR132 on macrophages

Thus far our work suggested that SEMA3D increases ARF6-GTP signaling to induce lactate secretion from PDA cells, which can reprogram macrophages. Next, we investigated the potential mechanism underlying macrophage reprogramming from secreted lactate by PDA cells. The G-protein coupled receptor 132 (GPCR132) has previously been shown to sense rising lactate levels in the acidic TME in breast cancer^40^. To examine if GPCR132 is involved in macrophage reprogramming due to SEMA3D signaling in PDA, M1 and M2 macrophages were co-cultured with KPC cells treated with SEMA3D-AP or CTRL-AP recombinant protein. We found that M1 and M2 polarized macrophages had significantly increased *GPCR132* mRNA expression of after co-culture with SEMA3D-AP treated KPC cells compared to CTRL-AP treated cells (Figure S5a and S5b). If GPCR132 is needed for SEMA3D signaling induced polarization changes, we hypothesized that BMDMs from a *GPCR132* KO mouse would lack changes in polarization after co-culture with SEMA3D-AP treated KPC cells. We observed that although polarization of BMDMs from WT and GPCR132 KO mice was similar as assessed by M1 and M2 gene expression markers (Figure S5c and S5d), the increase in gene expression markers in WT M2 macrophages that were co-cultured with SEMA3D-AP treated KPC cells (see Figure 3e) was no longer observed in M2 macrophages derived from *GPCR132* KO mice (Figure S5e and S5f). This data suggests that GPCR132 is required for SEMA3D macrophage reprogramming by PDA tumor cells.

### SEMA3D increases tumor-secreted lactate which impacts macrophage polarization in vivo

Since one of the sources of secreted SEMA3D in the stroma is tumor cells, which could affect lactate production in PDA tumor cells through a paracrine effect, we examined the effect of SEMA3D LOF in PDA cells on lactate production and macrophage reprogramming.

Immunohistochemical staining using F480, a macrophage marker, on PDA samples demonstrated that KPCS tumors have a significantly lower percentage of F480 strongly-positive cells compared to KPC tumors (Figure 4a and 4b). To examine the role of tumor cell-expressed SEMA3D on macrophage polarization in the TME, CD11b^+^ cells were isolated from KPC and KPCS dissected pancreas at ages of 12 and 16 weeks, respectively. M1 and M2 gene expression markers were analyzed in the isolated CD11b^+^ cells. At 12 weeks of age, CD11b^+^ cells isolated from the pancreas of KPCS mice exhibited a statistically significant reduction in the expression of some M2 markers as compared to age matched cells isolated from KPC mice (Figure 4c). Changes in M2 gene expression remained significant when CD11b^+^ cells were isolated from KPCS mice at 16 weeks of age compared to those from KPC mice. No difference was observed in M1 gene expression markers between KPC and KPCS mice, apart from KPCS mice expressing significantly less IL-6 at age of 16 weeks compared to KPC mice (Figure 4c). Therefore, the in vivo role of tumor cell-derived SEMA3D is consistent with the function of SEMA3D in reprogramming macrophages toward M2 polarization indirectly through tumor cells, as demonstrated by our in vitro co-culture experiments. SEMA3D LOF in mouse PDA cells skews the macrophages away from M2 polarization, accounting for the suppressive effects on PanIN progression, invasive tumor growth and, importantly, metastasis.

**Figure 4:**
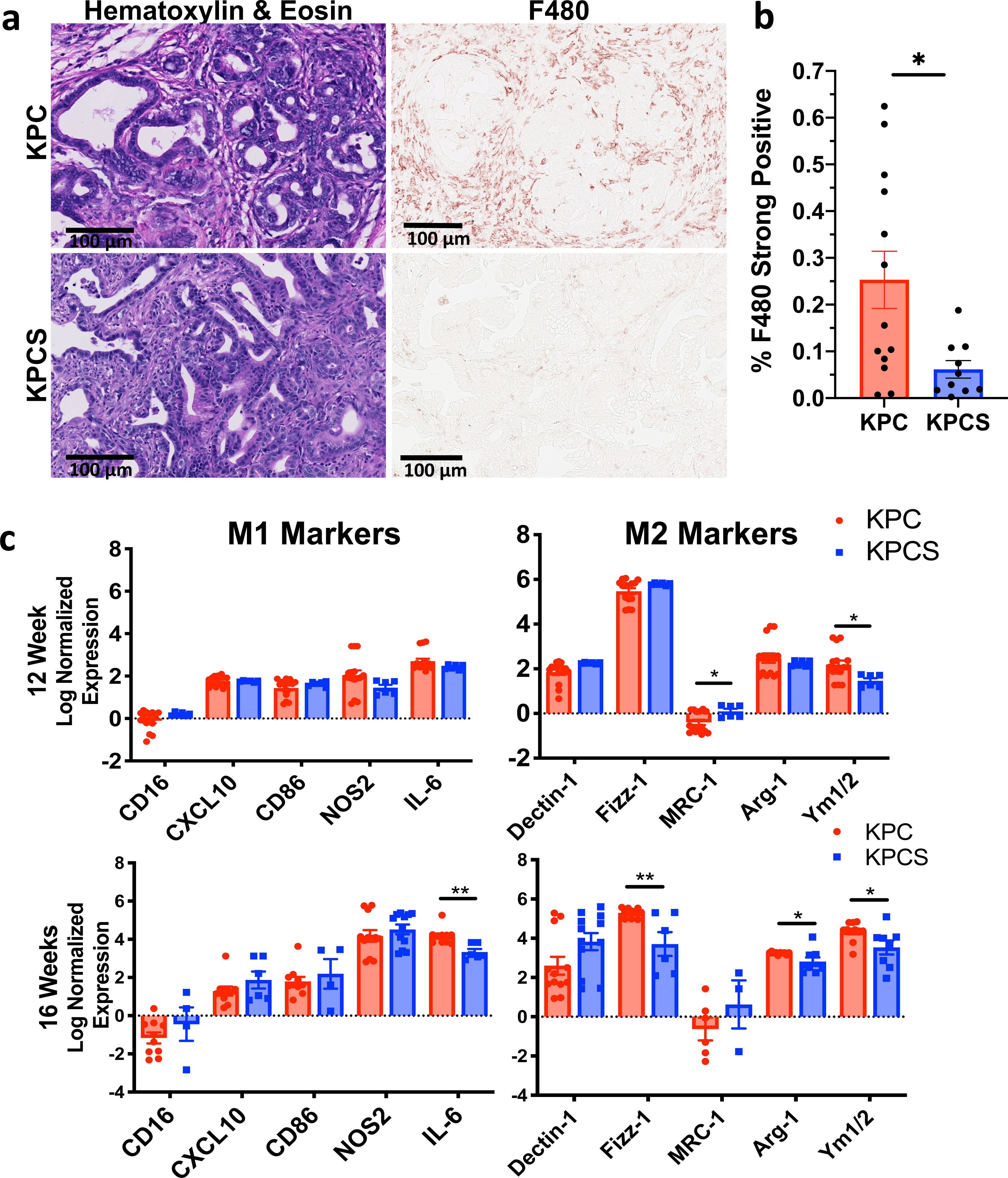
SEMA3D increases tumor secreted lactate which impacts macrophage polarization in vivo a. Representative images of immunohistochemical staining of F480 in KPC and KPCS PDA, respectively. b. Percentages of strongly positive, F480-stained cells in KPC (n=13) and KPCS (n=10) PDAs, respectively. c. qRT-PCR analysis of gene expression markers for M1 and M2 polarization in CD11b^+^ cells isolated from the pancreas of KPC and KPCS mice at age of 12 (top) and 16 (bottom) weeks, respectively.

### Nerve-derived SEMA3D impacts macrophage polarization

Another source of secreted SEMA3D in the stroma is from nerves in the TME. Therefore, we investigated whether or not SEMA3D secreted from nerves can contribute to increased lactate secretion by tumor cells and therefore impact polarization changes in macrophages. We previously observed that nerves within the TME in both murine KPC and human samples express SEMA3D and PLXND1^10^. To collect SEMA3D secreted from nerves, dorsal root ganglion (DRG) cells were grown in culture and DRG supernatant was collected and measured for SEMA3D protein levels; these were 203 pg/mL on average (Figure 5a). DRG-conditioned media was then used as another source of nerve-derived SEMA3D treatment on tumor cells. KPC cells treated with DRG-conditioned media also demonstrated increased ARF6-GTP protein expression compared to control media-treated cells (Figure 5b). Furthermore, pretreatment of KPC cells with the PLXND1 neutralizing antibody before exposure to DRG-conditioned media inhibited the increased expression of ARF6-GTP, indicating DRG-conditioned media activates ARF6-GTP in a PLXND1- dependent manner; this is similar to what we observed using SEMA3D-AP recombinant protein treatment (Figure 5c). The addition of DRG-conditioned media to KPC cells also induced a significantly higher mRNA expression of the glycolysis genes *GLUT-1*, *HK-2* and *LDHA* following IgG isotype treatment, and this was diminished upon PLXND1 neutralizing antibody treatment (Figure 5d). Interestingly, DRG-conditioned media treatment induced higher *GLUT-1, HK-2* and *LDHA* mRNA expression compared to recombinant SEMA3D-AP treatment alone, suggesting that another semaphorin or other signaling molecule also signals through PLXND1 to increase glycolysis in KPC cells (Figure 5d).

**Figure 5:**
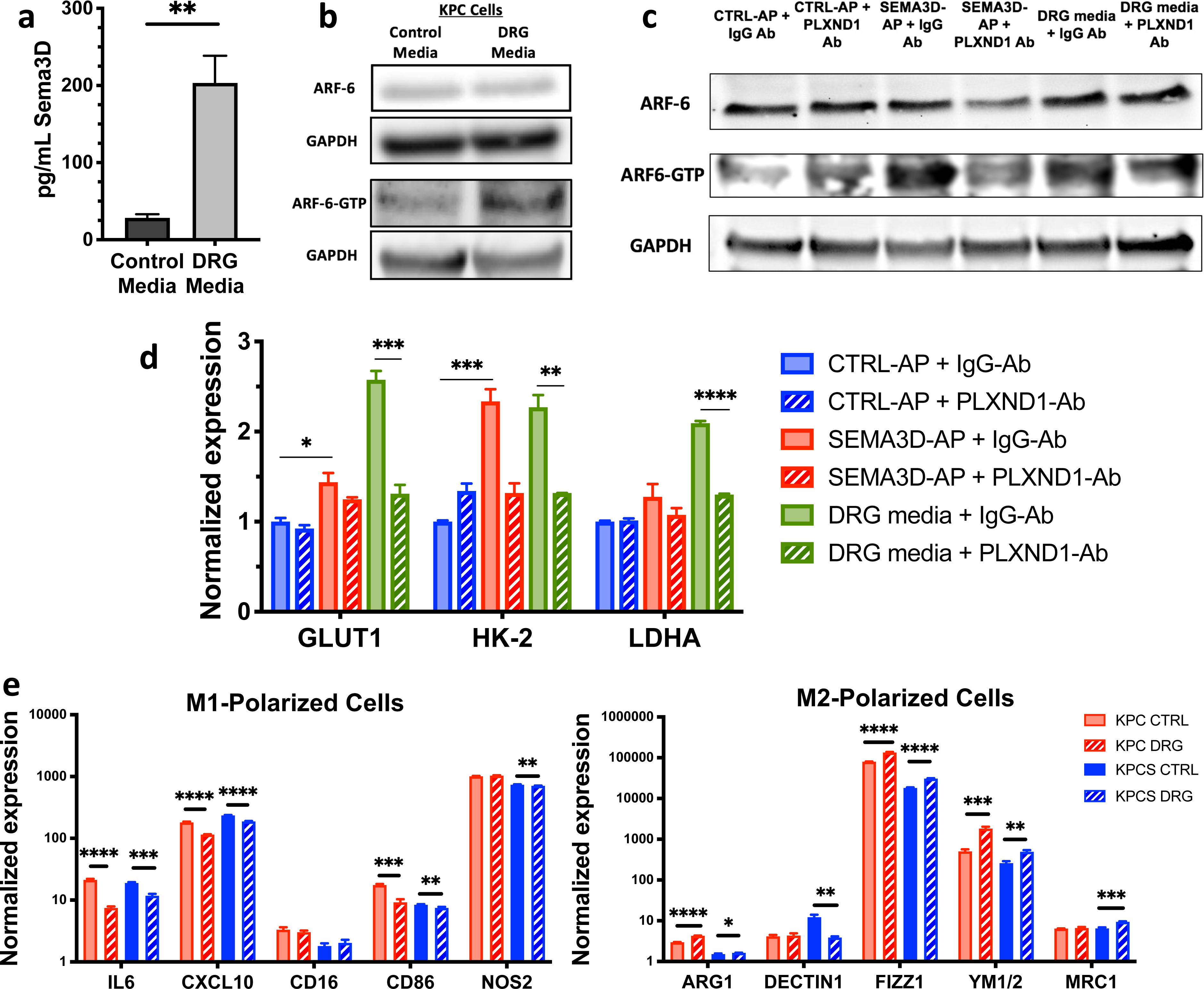
Nerve-derived SEMA3D impacts macrophage polarization in vitro a. ELISA analysis of secreted SEMA3D was performed on media cultured with and without DRG cells. b. Western blot analysis of the expression of ARF6 and active ARF6-GTP proteins in KPC cells treated with control and DRG-conditioning media, respectively. c. Western blot analysis of ARF6 and active ARF6-GTP proteins in KPC cells treated with CTRL-AP, SEMA3D-AP or the DRG- conditioning media supplemented with anti-PLXND1 neutralizing antibody or isotype IgG control. d. qRT-PCR analysis of gene expression of *GLUT-1*, *HK-2*, and *LDHA* in KPC cells treated with CTRL-AP, SEMA3D-AP or the DRG-conditioning media supplemented with anti-PLXND1 neutralizing antibody or isotype IgG control. e. qRT-PCR analysis of gene expression markers for M1 and M2 polarization in M1- and M2- polarized macrophages, as indicated, after co-culture with KPC or KPCS cells treated with control or DRG-conditioning media.

Since DRG-conditioned media is able to induce the activation of ARF6-GTP and glycolytic gene expression in tumor cells, we next analyzed the polarization of macrophages after co-culture with tumor cells exposed to DRG-conditioned media. KPC and KPCS cells were used for BMDM co-culture to examine cells with and without endogenous SEMA3D expression. Several M1 gene expression markers were decreased in M1-polarized macrophages following exposure to either KPC or KPCS cells co-cultured with DRG-conditioning media compared to KPC or KPCS cells co-cultured with control media (Figure 5e). M2-polarized macrophages co-cultured with either KPC or KPCS cells treated with DRG-conditioned media also displayed an increase in most M2 gene expression markers compared to control media-treated KPC or KPCS co-culture, respectively (Figure 5e). Taken together, these results suggest that nerve-derived SEMA3D signaling through PLXND1 activates ARF6-GTP in tumor cells and induces the expression of glycolytic mRNAs to skew macrophages toward M2 polarization, regardless the status of endogenous SEMA3D. These results also suggest that nerve-derived SEMA3D can reprogram macrophages through activating ARF6-GTP and subsequently by enhancing glycolytic gene expression in PDA.

### SEMA3D-induced ARF6-GTP signaling is mutant KRAS-dependent

Previously, SEMA3E binding to PLXND1 was observed to activate the R-RAS signaling^34^. ARF6 is a well-characterized downstream target of KRAS^38^. The Ras signaling pathway is of particular importance in the context of PDA since almost 90% of patients have a GOF mutation in the KRAS gene, and the mutated KRAS protein and its downstream signaling cascade have proven to be difficult with respect to drug targeting^41,42^. Oncogenic KRAS signaling activates ERK1/2, which subsequently activates ARF6-GDP to ARF-GTP to increase glucose metabolism in cancerous cells^38^. To examine if SEMA3D-induced ARF6-GTP signaling is dependent on KRAS mutations in KPC cells, we next examined the impact of SEMA3D treatment on a cell line without a GOF mutation in KRAS. Panc02 cells were generated from a 3-methy-cholanthrene-induced pancreatic tumor cell line derived from C57Bl/6 mice that express wild-type KRAS^43^. Panc02 cells treated with SEMA3D-AP do not exhibit a significant increase in ARF6-GTP compared to CTRL-AP treated cells (Figure 6a). Interestingly, Panc02 cells did display an increase in the levels of the non-active form of the ARF6, ARF6-GDP protein when treated with SEMA3D-AP (Figure 6a).

**Figure 6:**
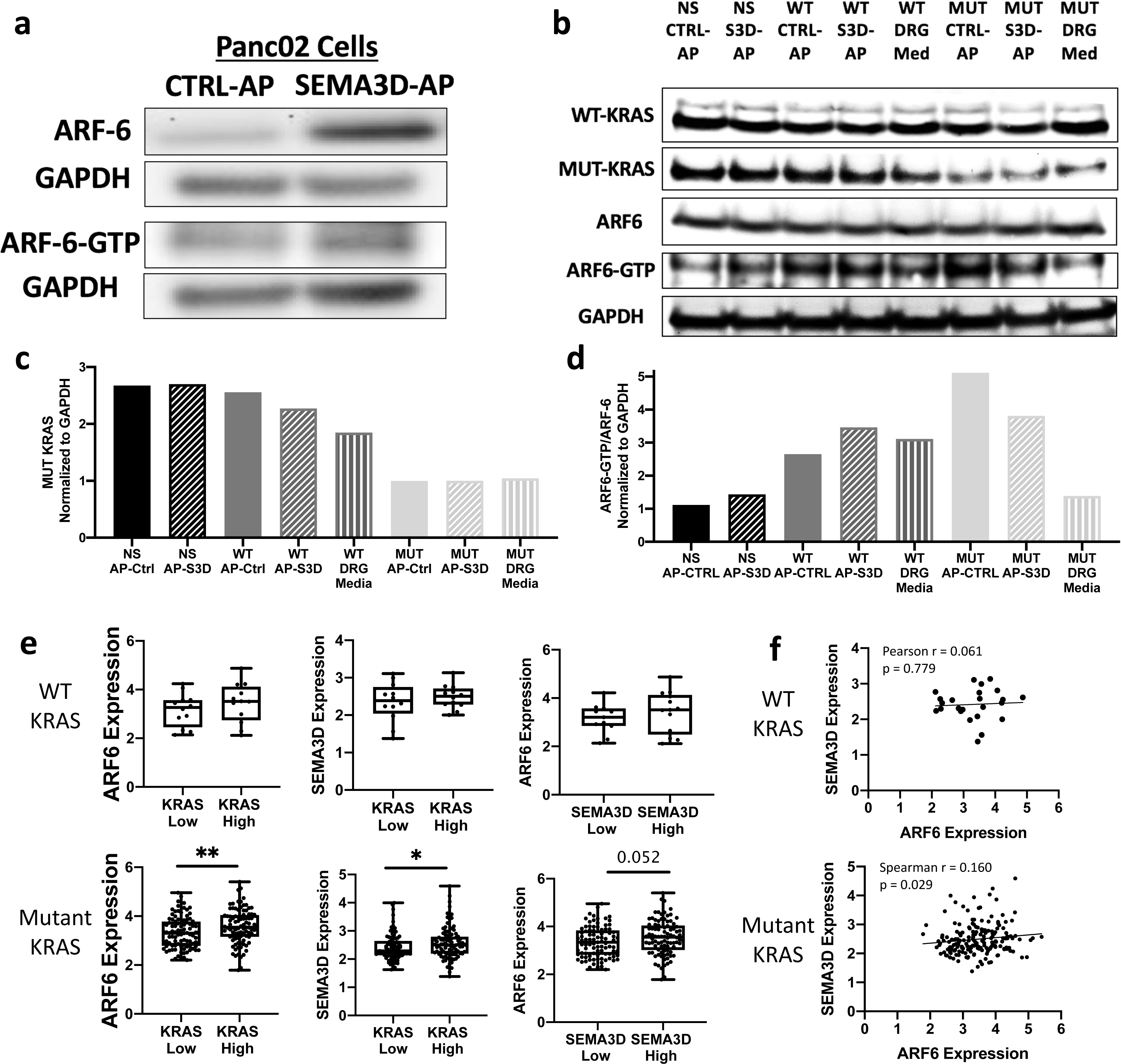
Mutated KRAS-dependency of *SEMA3D-induced ARF6-GTP signaling* a. Western blot analysis of ARF6 and active ARF6-GTP protein expression in Panc02 cells treated with CTRL-AP and SEMA3D-AP recombinant protein, respectively. b. Western blot analysis of WT and MUT KRAS proteins as well as ARF and ARF-GTP proteins in non-specific (NS), WT, or mutant (MUT) siRNA-treated KPC cells that were cultured with either the CTRL-AP or SEMA3D- AP containing media or the DRG-conditioning media. c. Densitometry quantification of MUT KRAS protein normalized to GAPDH according to the Western blot analysis (b). d. Densitometry quantification of ARF6-GTP/ARF6 protein expression ratio normalized to GAPDH according to the Western blot analysis (b). e. KRAS, ARF6 and SEMA3D gene expression analysis using normalized RNA sequencing expression data derived from 209 human PDA samples annotated with the KRAS WT (top, n=24) or MUT (bottom, n=185) status. WT or MUT KRAS PDA samples were subgrouped into KRAS high vs. low or SEMA3D high vs. low, respectively, by using their median expression value as the cut-off. f. Scatter plots of normalized gene expression levels of SEMA3D and ARF6 in KRAS WT or MUT human PDAs, respectively.

Next, KPC cells with *KRAS^G12D^*-specific siRNA knockdown^44^ were created to further examine the effect of SEMA3D treatment on KPC cells with reduced mutant KRAS signaling. KRAS was knocked down in KPC cells using siRNA targeting *KRAS^G12D^*, *KRAS^WT^*, and a non-specific (NS) control. Knockdown was confirmed using KRAS^G12D^ and KRAS^WT^ specific antibodies to detect protein expression (Figure 6b and 6c). After siRNA knock-down, the KPC cells were treated with CTRL-AP, SEMA3D-AP or DRG-conditioning media. In the NS siRNA- or *KRAS^WT^* siRNA-treated KPC cells, in which the mutant *KRAS^G12D^* is still expressed, SEMA3D-AP treatment increased ARF6-GTP protein levels compared to CTRL-AP treatment. However, no increase in ARF6-GTP protein was observed in the *KRAS^G12D^* siRNA-treated KPC cells treated with SEMA3D or DRG-conditioned media (Figure 6b and 6d). These results suggest that SEMA3D-induced ARF6-GTP signaling is dependent upon the presence of mutated KRAS*^G12D^*.

To examine the impact of human KRAS mutations on SEMA3D-induced ARF6 signaling, Illumina RNA sequencing data from the ICGC database were examined to compare gene expression in 209 PDA samples from patients previously identified as expressing either KRAS^WT^ (n=185) or KRAS^MUT^ (n=185) PDA tumors^45^. Samples were separated into KRAS^high^ and KRAS^low^ expressing groups by using the median value of KRAS expression as a cut-off. PDA samples expressing high levels of

KRAS^MUT^ were found to express significantly higher levels of ARF6 compared to samples with low KRAS expression, aligning with previously published results^38^ (Figure 6e). This increase in ARF6 expression was not observed in samples expressing higher levels of KRAS^WT^. Furthermore, PDA samples expressing high KRAS^MUT^ (KRAS^high^) levels had significantly higher expression levels of SEMA3D compared to KRAS^low^-expressing samples. Similarly, this correlation was not observed in samples with KRAS^WT^ (Figure 6e). In addition, samples were separated into SEMA3D^high^ and SEMA3D^low^ expressing groups by using the median value of SEMA3D expression as the cut-off. Samples expressing KRAS^MUT^ and high levels of SEMA3D trended toward having higher levels of ARF6 expression compared to SEMA3D^low^ expressing samples. No difference in ARF6 expression was observed between SEMA3D^high^ and SEMA3D^low^ expressing KRAS^WT^ samples. Moreover, in PDA samples expressing KRAS^MUT^, a significant positive correlation was found between ARF6 and SEMA3D expression (Spearman r = 0.160, p value = 0.029); in contrast, no significant correlation was found with patients expressing KRAS^WT^ (Figure 6f). These results suggest that higher KRAS^MUT^ expression is associated with an increase in SEMA3D and ARF6 expression in human PDAs. Taken together, these data support the correlation involving increases in KRAS^MUT^, ARF6-GTP, and SEMA3D expression in PDAs.

### Increased number of M2 macrophages proximal to nerves in human PDA expressing SEMA3D

To further investigate the role of tumor and nerve secreted SEMA3D on macrophage polarization, human PDA samples were evaluated via multiplex immunohistochemical analysis as previously described^10,46^. Human PDA samples were stained for TUJ1 to identify nerves within the TME, which acts as an additional source of SEMA3D. Three unique regions of interest were identified in proximal (within 650mm) or distal (greater than 650mm to a nerve bundle) regions, respectively, in each human PDA sample. Immunohistochemical staining of CD45, CSF-1R, CD163, and CD68 was used to identify M1 and M2-like tumor-associated macrophages (TAMs). Aligning with previous work^47^, our results show that M2-like TAMs constitute a higher proportion of the CD45^+^ cells in the human PDA TME, whereas only a small percentage of CD45^+^ cells was M1-like TAMs (Figure 7a). Furthermore, there was no significant difference between the percentages of M1-like TAMs either proximal or distal to nerves in human PDAs. In contrast, a significant increase in M2 TAM percentage was observed in areas of the TME proximal to nerves compared to areas distal to the nerves (Figure 7a). These results suggest that the intratumoral nerves skew nearby macrophages toward M2 polarization, supporting a role for nerves, or factors-secreted by neurons, in macrophage reprogramming.

**Figure 7:**
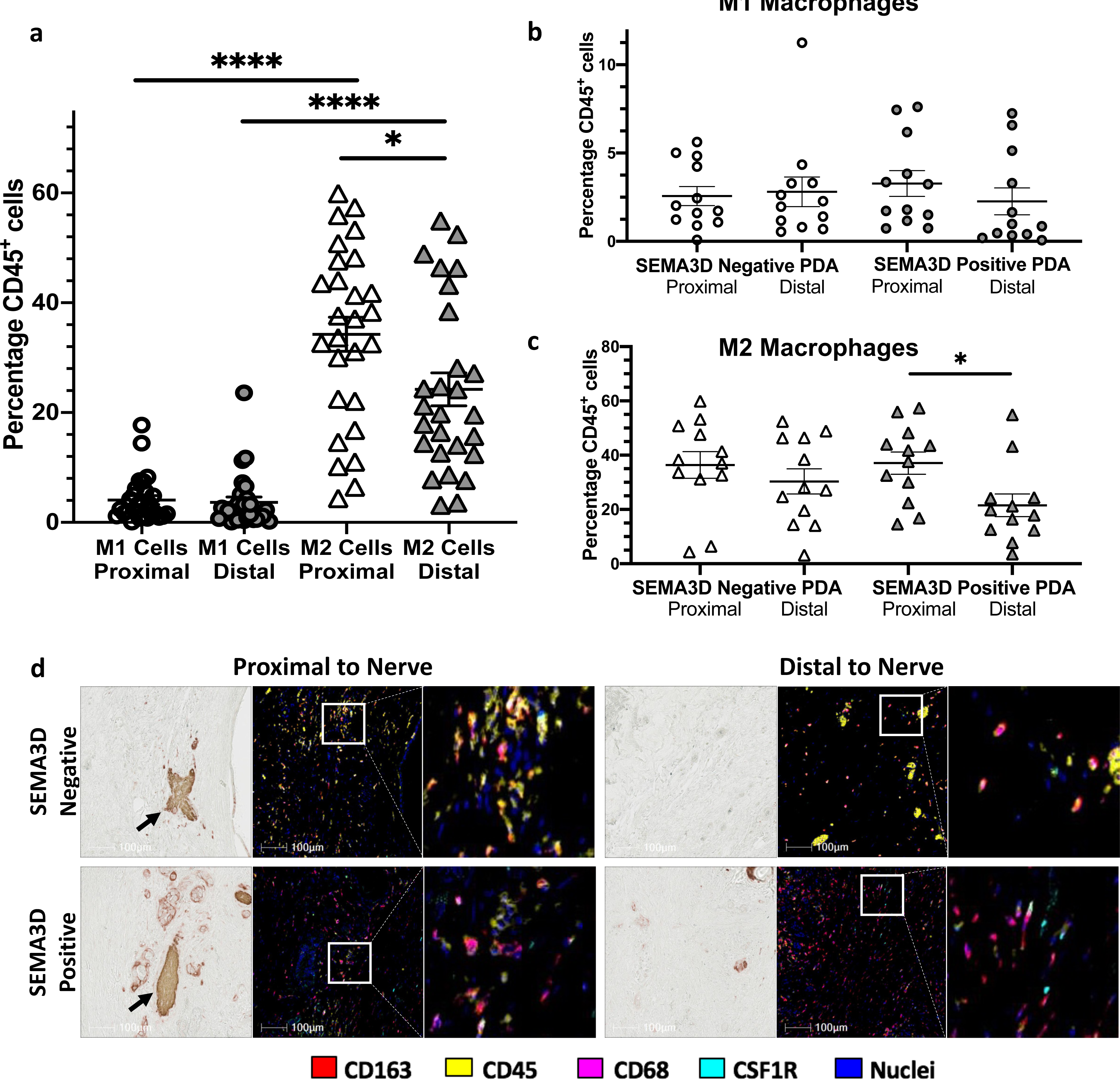
Increase in M2 macrophages proximal to nerves in human PDA expressing SEMA3D a. Percentages of M1 or M2-like TAMs distal or proximal to nerves in human PDA. b,c. Percentages of M1(b) or M2(c)-like TAMs distal or proximal to nerves in SEMA3D-positive and - negative human PDA samples. d. Representative images of multiplex immunohistochemical staining of SEMA3D-negative and -positive human PDA samples in areas proximal or distal to nerves, respectively. Nerves identified with anti-TUJ1 staining (arrows) and immune cells identified with anti-CD45, CD163, CSF1R, and CD68 staining. An amplified region in each panel was shown (white box).

To address this issue further, patient samples were separated into SEMA3D negative or SEMA3D positive samples on basis of immunohistochemical staining for SEMA3D on tumor epithelial cells, as previously described^10^. M1-like TAMs were found in similar fractions proximal or distal to nerves in both SEMA3D-positive and -negative PDA samples, respectively (Figure 7b). However, in SEMA3D-positive PDA samples, M2 TAMs were observed to be present in higher percentages proximal to nerves as compared to those distal to the nerve in the PDA TME (Figure 7c). These results suggest that both SEMA3D expression in the PDA tumor cells and in the nerves within the vicinity of tumor cells together play a role in reprogramming TAMs toward procancerous M2 polarization. Since our previous work demonstrated the association of SEMA3D with PNI, our new finding provides a mechanism for understanding the association between PNI and poor prognosis in PDA.

## DISCUSSION

In this study we present one of the first demonstrations of a mechanistic role for axon guidance molecules in PDA development and metastasis. Previously, genes encoding axon guidance molecules have been observed to be some of the most frequently affected with respect to mutations and copy number changes in human PDA. Our previous work revealed a mechanistic role for SEMA3D in mediating the interaction between tumor cells and nerves in PDA. However, this study provides the first description of the mechanism underlying how this interaction promotes PDA development and metastasis. We demonstrate here both the role of tumor-derived SEMA3D in the malignant transformation of pancreatic epithelial cells and also the role of nerve-derived SEMA3D in PDA progression. We also reveal for the first time the change in TAM polarization as a mechanistic role of tumor-intrinsic SEMA3D. These results also provide new evidence at the molecular level supporting previously characterized autocrine and paracrine functions of SEMA3D in PDA progression and metastasis^10,15^. Interestingly, nerve-derived SEMA3D does not modulate TAMs directly, but does so indirectly through tumor cells. On one hand, we reveal a compensatory mechanism dependent on tumor-extrinsic SEMA3D. On the other hand, our work suggests that the same mechanism may apply to both tumor-intrinsic and -extrinsic SEMA3D, and that this is mediated through lactate produced by tumor cells as a result of the Warburg effect, which in turn reprograms macrophages.

For the first time, SEMA3D is also shown to induce lactate production and subsequent macrophage reprogramming in a mutated KRAS-dependent manner. Our study also shows a normal function of axon guidance molecules in the nerves, and not just their atopic expression in cancer cells^13–15,19,20,48–50^, can contribute to cancer progression and metastasis through an oncogenic KRAS-ARF6 pathway. Together, our work demonstrates how inflammatory responses in the TME promote PanIN and invasive PDA progression and metastasis driven by an oncogenic process (see proposed model in Figure 8). Further elucidation of axon guidance signaling pathways in the PDA TME will provide new therapeutic targets to prevent PNI and metastatic progression of PDA and improve overall survival of patients.

**Figure 8:**
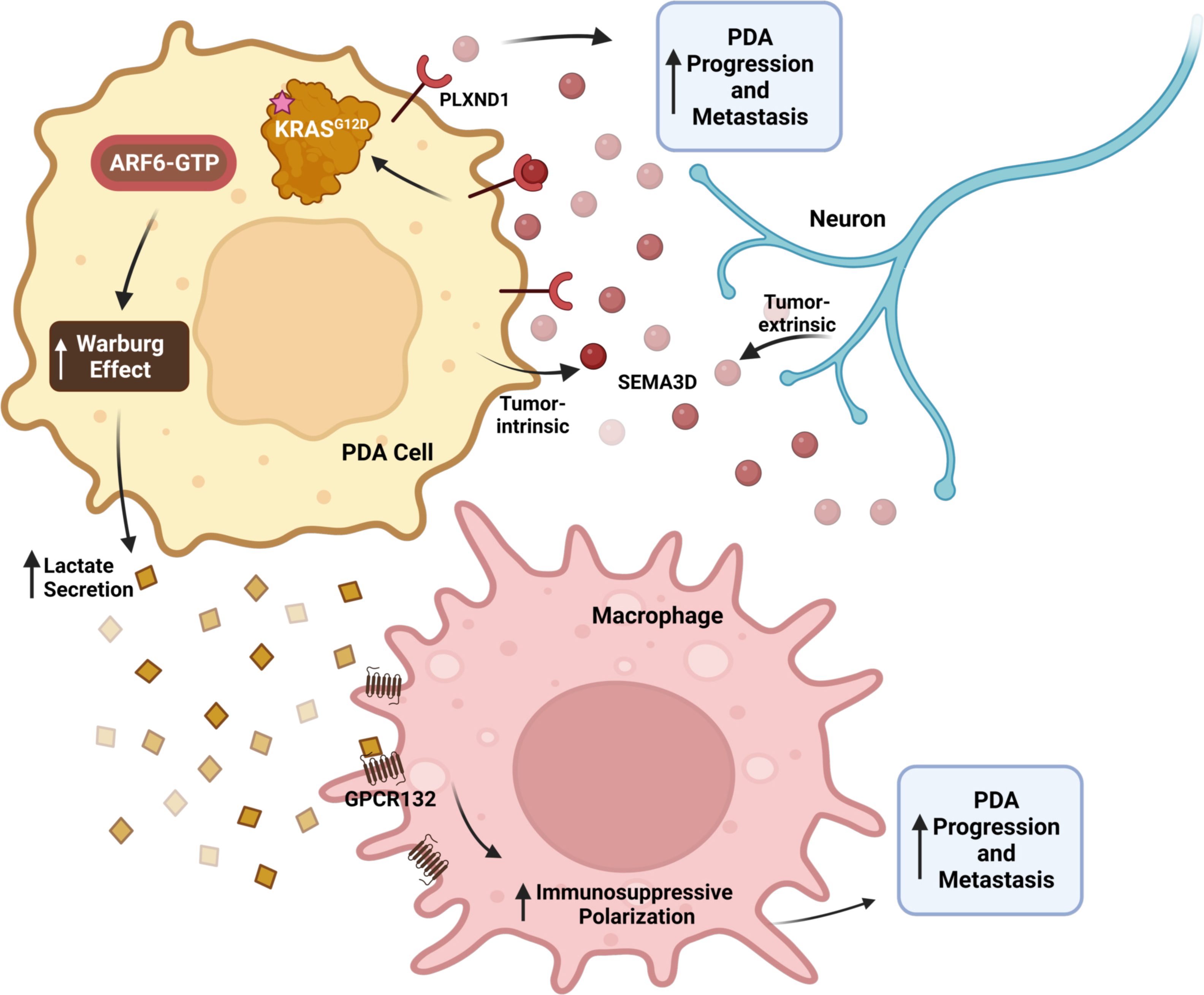
Tumor-intrinsic and -extrinsic SEMA3D signaling reprograms macrophages to aid PDA progression and metastasis Both tumor-intrinsic SEMA3D and tumor-extrinsic, neuron-derived SEMA3D promote PDA progression and metastasis. Autocrine or paracrine SEMA3D binds to its receptor, PLXND1, on PDA cells. The SEMA3D-PLXND1 signaling is mediated by mutant KRAS to activate ARF6-GTP and subsequently increase glycolytic gene expression. Increased production of lactate, as a result of Warburg effect, is sensed by GPCR132 on macrophages, which are subsequently reprogrammed to become M2 polarized. The protumoral M2 macrophages promote PDA progression and metastasis. Created with Biorender.com.

We also developed the KPCS transgenic mouse to examine the role of tumoral SEMA3D throughout the stepwise progression of PDA. Creation of a model with global loss of SEMA3D was not possible as global SEMA3D knockout mice only survive a few hours after birth because newly born mice cannot be breast fed due to the loss of taste sensation. Interestingly, we found that KPCS mice did not develop any metastasis in the lung, liver, gut or peritoneum, supporting a role of SEMA3D in metastasis formation. However, this role of SEMA3D may be restricted to tumor-intrinsic SEMA3D. It is possible that nerve-derived SEMA3D could provide compensation for metastasis formation, but mice do not survive the burden of the primary tumor and die before the development of metastasis. Supporting this notion is the observation of PNI in the primary tumor of KPCS mice. Nevertheless, only a modest reduction in PNI and nerve density in primary tumors from KPCS mice was found, suggesting that other axon guidance pathways may serve to compensate and remain to be explored. Consistently, multiple axon guidance family members have had genetic alterations in individual human PDAs^11^. Our study demonstrated that SEMA3D expression correlates with TAM infiltration and polarization in the TME of human PDAs. It will also be interesting to examine whether expression and genetic alteration of other axon guidance and neuronal development genes would have an impact on the immune cell infiltrates and have a similar dual role in human PDAs. Furthermore, a recent study found neurons in locations remote to tumors can promote malignant progression of cancer, which also remains to be explored in PDA^51^. Therefore, we do not think that a single axon guidance molecule itself, but a combination of multiple axon guidance family members together would impact the overall prognosis.

Published studies have established the role of TAMs in the malignant transformation of pancreatic epithelial cells^28–30,47,52–54^. The role of oncogenic KRAS in regulating anabolic glucose metabolism in PDA is also well established^55^. However, TAM reprogramming has not been linked to the intratumoral KRAS-ARF6 pathway or to the axon guidance molecules on the nerves. Interestingly, we did not identify a direct role for SEMA3D in macrophages due to lack of the SEMA3D receptor expression. Although a direct role for other axon guidance molecules in modulating macrophage functions cannot be ruled out, our study demonstrated that nerve-derived SEMA3D reprograms macrophages indirectly through tumor cells. This suggests that nerve-derived factors do not naturally modulate the TME, but tumor cells “hijack” the nerves signaling to reprogram the TME and skew it toward one that favors cancer progression and metastasis. It will be interesting to investigate whether the genetic alterations in the axon guidance molecule receptors, such as plexins on tumor cells, would have an impact on this mechanism.

Thus, this study has expanded “Cancer Neuroscience” into a new area^56^. Cancer cells not only acquire neuronal cell features, but also “hijack” the nerves to help them reprogram their TME into one that favors their growth, progression and metastasis. This study also shows that SEMA3D is a potential target for cancer therapeutics. As both tumoral and nerve-derived SEMA3D play a role in PDA progression and metastasis, targeting all sources of SEMA3D would be critical as a therapeutic target. Therefore, an antibody TRAP platform may be an attractive drug development strategy for SEMA3D. Future studies will be focused on developing therapeutic agents that target SEMA3D and also on revealing how multiple axon guidance molecules coordinate together in promoting PDA progression and metastasis.

## METHODS

### Material Availability

Mouse lines generated in this study will be shared upon contact with the lead contact.

### Data and Code Availability

All data reported in this paper and any additional information required to reanalyze the data reported in this paper is available from the lead contact upon request. This paper does not report original code.

## EXPERIMENTAL MODELS AND SUBJECT DETAILS

### Mouse Models of PDA

All animal experiments were executed in accordance with the Animal Care and Use Committee of Johns Hopkins University and animals were maintained according to the American Association of Laboratory Animal Care guidelines. The *LSL-KRAS^G12D/+;^LSL-TRP53^R^*^172^*^H/+^;PDX-1-Cre^+/+^* (KPC) mouse is a genetically engineered mouse model of PDA, previously established though a pancreatic specific knock-in of conditional alleles of the KRAS^G12D^ and TP53^R172H^ mutations on a mixed background of 129/SvJae/C57Bl/6^32^. When crossed with PDX-1-CRE^+/+^ mice, who express pancreas-specific cre recombinase, they develop PanIN lesions that progress stepwise to full PDA development similar to human disease. Mice with conditional expression of SEMA3D, in which SEMA3D is flanked by loxP sites (SEMA3D^flox/flox^), was obtained from personal communication with Dr. David Ginty. When crossed with KPC mice, the result is LSL-KRAS^G12D/+^; LSL-TRP53^R172H/+^; LSL-SEMA3D^flox/flox^; PDX-1-Cre mice, or KPCS mice with SEMA3D and TP53 loss of function and KRAS gain of function, specifically in the pancreas. Mice genotyping was confirmed through Transnetyx. Both male and female mice were examined at indicated time-points. Tumor sizes of KPC and KPCS mice were monitored at the indicated time points using small animal ultrasound technologies (Vevo770, Visual Sonics).

### Murine Cell Culture

KPC and KPCS cell lines were developed as previously described from primary tumors of KPC and KPCS mice respectively^15^. The Panc02 cell line was generated from a 3-methy-cholanthrene- induced pancreatic tumor cell line derived from C57Bl/6 mice^43^. The KPC, KPCS and Panc02 cells were cultured in RPMI 1640 medium supplemented with 10% FBS, 1mM sodium pyruvate, 2mM L-glutamine, 1% nonessential amino acids (100x), 50 units/mL penicillin and 50 units/mL streptomycin (Invitrogen) at 37°C in 5% and 10% CO_2_, respectively. COS7 and primary DRG cells were cultured in Dulbecco’s Modified Eagles Medium supplemented with 10% FBS at 37°C in 5% CO_2_. Neutralizing antibody against goat anti-PlexinD1 or goat IgG isotype control (Novus) was used during cell culture.

Primary DRG cells were collected and cultured from postnatal day 1-7 mice as previously described^10^. In summary, mice were euthanized and the entire vertebral column was transferred to Leibovitz’s L15 medium (Gibco) in a 100mm cell culture dish with the dorsal side facing up under a microscope (Nikon SMZ Stereo Zoom). The vertebral column was cut and connections between the DRG and spinal cord were severed and the spinal cord was removed. DRGs were pulled out individually and kept in Leibovitz’s L15 medium on ice. After collection, DRGs were spun down at low speed and resuspended in 5mg/mL collagenase and 2mg/mL dispase and incubated for 40 minutes shaking at 37°C. Cells were then washed and cultured in Dulbecco’s Modified Eagles medium supplemented with 10% FBS at 37°C in 5% CO_2_.

## METHOD DETAIL

### Western Blot Analysis

Cells were lysed in IP lysis buffer (Thermo Scientific) containing protease and phosphatase inhibitors. After lysis, the lysate was spun at 15,000 rpm for 5 minutes and the samples were boiled in SDS containing reducing agent (Bio-Rad). The samples were loaded and electrophoresed on a 4-12% bis-tris gel (Bio-Rad) for 2 hours at 100 V. The gels were transferred onto nitrocellulouse membranes for 1 hour at 80V. The membranes were blocked in 5% bovine serum albumin (BSA) for 1 hour on a shaker at room temperature. Primary antibodies were added in 2.5% BSA overnight on a shaker at 4°C. The membranes were washed with 1xTBST and then incubated with rabbit or mouse secondary antibodies against horseradish peroxidase (1:5000; GE) for 30 minutes to 1 hour at room temperature. The membranes were developed using enhanced chemiluminescent reagents (GE).

### Quantitative RT-PCR

RNA was isolated from cells using Trizol reagent according to manufacturer protocol. In summary, the cells were incubated in 1 ml of Trizol reagent at room temperature for 5 minutes. Chloroform was added and the samples were shaken vigorously for 15 s before incubation at room temperature for two minutes. Samples were spun and the aqueous phase was transferred to a fresh microcentrifuge tube. Isopropanol was added to the aqueous phase, and the samples were left at room temperature for ten minutes. The samples were again centrifuged at 12,000 rpm and the RNA pellet was washed once in 75% ethanol, then centrifuged at 9500 rpm, and left to air-dry for 30 minutes at room temperature. The RNA pellet was resuspended in 50 ul of distilled water and the concentration was measured by Nanodrop. Reverse transcription was performed using RDRT ReadyScript cDNA synthesis Mix (Sigma-Aldrich). qRT-PCR was performed on an Applied Biosystems RT-PCR machine (Life Technologies). All primers were obtained from Invitrogen. Reactions were performed using SYBR Green PCR master mix (Applied Biosystems). All mRNA expression values were normalized to mouse *BETA-ACTIN* expression values. Polarized macrophage expression was normalized against non-polarized M0 macrophages.

### Immunohistochemistry

Immunohistochemistry on mouse tissue was performed manually with Anti-Rabbit IgG ImmPRESS Excel Staining Kit (Vector). After deparaffinization and rehydration of the tissue, heat-induced antigen retrieval for SEMA3D staining was performed in EDTA buffer (pH 9.0) using a pressure cooker at 125°C for 30 seconds and 95°C for 10 seconds. Slides were placed in Bloxall blocking solution for 10 minutes, washed in 1xTBST for 5 minutes, and incubated with kit supplied horse serum (2.5%) for 20 minutes. Incubation with rabbit antibodies against SEMA3D (Abcam) at 1:200 for 1 hour was followed by 2 1xTBST washes and kit supplied secondary amplifier antibody for 1 hour. After washing, ImmPRESS Excel polymer reagent solution was then added for 30 minutes and washed again. 3,3’-Diaminobenzidine hydrochloride was added for development. All slides were counterstained with hematoxylin. Stained slides were scanned using an Aperio ScanScope AT (Leica Biosystems) at 40x magnification.

Anti-F480 antibody was used to stain mouse macrophages as described previously^33^. Percentage of strong positive F480 staining was determined as the number of the strongly positive stained cells divided by the number of total cells in the specimen area using the Positive Pixel Count 2004- 08-11 algorithm version 8.100 (Aperio Technologies)^57^. Four specimen areas were measured for percent of strong positive cells in each mouse sample and the results were averaged as previously described^57^.

Nerves in the tumor area were analyzed by TUJ1 immunohistochemical staining and analyzed based on positive stained cells over total cells using Aperio ImageScope.

### Generation of Recombinant AP-Sema3D and AP-Control

Production of recombinant AP-SEMA3D and AP-Control was produced as described previously^15^. Briefly, COS7 cells were seeded into a 100mm dish at 80% confluency. Cells were transfected using 12 µg of the plasmid DNA (AP-SEMA3D or AP-Control) with Lipofectamine 2000 (Invitrogen) in serum media according to the manufacturer’s protocol. Twenty-four hours later, the media was replaced with serum-free media. Forty-eight hours later, the media was collected and filtered using a 0.22µm syringe for subsequent use.

### RNA interference

To perform RNA interference cells were seeded at 80% confluence in a 100mm dish. Eighty pmol of WT KRAS, G12D KRAS and control scramble siGENOME siRNA (GE) was transfected with Lipofectamine 2000 in serum media according to the manufacturer’s protocol (Invitrogen). Twenty-four hours after transfection the culture media was replaced with serum-free media. Forty-eight hours after transfection, the cells were treated with AP-CTRL, AP-SEMA3D or DRG conditioned medium.

### Macrophage Polarization

The bone-marrow derived macrophages (BMDMs) were isolated from healthy C57BL/6 or GPCR132 knock-out mice^40^. BMDMs were harvested with some modification according to previously described method^40,58^. Briefly, mouse hind legs were cut at the hip joint and excess muscle was removed from legs. The femur was severed proximal to each joint, and the bone cavity was flushed with at least 5 mL of transfer medium (RPMI 1640, 5% FBS). BMDMs were plated at 1 × 10^6^ cells per 10 cm dish (Corning Lifesciences, Tewksbury, MA, USA) with 10 mL macrophage complete medium (RPMI 1640, 20% FBS), containing 50ng/mL mouse recombinant macrophage colony-stimulating factor (M-CSF; Biolegend) for 7 days, with changes of media and M-CSF at day 5. Mouse BMDMs were treated with 400 ng/mL LPS (Sigma) and 30 ng/mL IFN-γ (Biolegend) for 24 hours to induce the M1 polarization at day 6; while, M2 polarization was induced through the addition of 50 ng/mL IL-4 (Biolegend) at day 6. During lactate induced polarization of BMDMs, 25 nM lactate was added for 24 hours to BMDM 6 days after harvest. KPC co-culture, CTRL-AP, SEMA3D-AP, or DRG media treatment was added 24 hours after polarization for 48 hours. KPC co-culture with BMDMs was performed using a transwell system in which BMDMs were placed on the bottom well and tumor cells on the top well. The chambers were separated by 1-micron pores to allow the passage of secreted factors and media to both cell types.

### CD11b^+^ Isolation from KPC/KPCS Mice

The pancreas was harvested from KPC and KPCS at indicated time-points and stored in RPMI with 5% FBS on ice during collection. The pancreas was cut into small 2-4 mm pieces in a 10-cm dish and transferred into a gentleMACs C tube containing tumor dissociation digestion mix (Militenyi Biotec). The tissue was dissociated using the gentleMACs dissociator. After dissociation, the tissue was spun and the sample was resuspended in macrophage media (RPMI 1640, 20% FBS) and run through 70μm strainer to create single cell suspension. Next, ACK lysis buffer was added for 1 minute to lyse red blood cells and was immediately quenched with macrophage media. Next the buffy coat was isolated from the cells using percoll. Cells were resuspended in 80% percoll and overlayed with 40% percoll and subsequently spun at 3200 rpm for 25 minutes without the brake. The buffy coat layer was isolated and quenched with macrophage media. Next the cells were resuspended with the positive cell collection CD11b microbead kit (Militenyi Biotec) per manufacturers instruction. The bead-cell mixture was run through an MS column with MACs buffer (Militenyi Biotec) and positively selected cells were eluted by removing the MS column from the magnet. The cells were then used to proceed with the quantitative RT-PCR protocol described above.

### Sema3D ELISA

Concentrations of mouse SEMA3D was determined using SEMA3D ELISA kit (Cusabio Biotech Co) per manufacturers instruction. In brief, supernatant from KPC, KPCS and DRG cells were collected after 48 hours. Supernatants were diluted in sample buffer and absorbance was read at 450 nm.

### Lactate Assay

The L-Lactate Assay kit (Abcam) was used per manufacture instructions. Media from KPC cells was collected after 48-hour treatment of CTRL-AP or SEMA3D-AP conditioned media. The kit utilizes the oxidation of lactate by lactate dehydrogenase to generate a product which interacts with a probe to produce a color that can be detected at 450 nm.

### ICGC Database Analysis

Previously published Illumina whole exome sequencing libraries for 209 human PDA samples were accessed from the International Cancer Genome Consortium (ICGC) and aligned with previously reported characterization of KRAS wild-type (n=24) or mutant status (n=185)^45^. The patient illumina sequencing normalized expression data was extracted for *KRAS*, *SEMA3D*, and *ARF6* genes. KRAS or SEMA3D high and low expressing samples were separated in two groups using the median expression value of KRAS or SEMA3D respectively.

### KPC and KPCS Mice Histology

The pancreas from KPC and KPCS mice was harvested at indicated time-points. The pancreas was fixed in formalin and paraffin embedded by the JHU reference laboratory. The formalin fixed paraffin embedded (FFPE) samples were then cut and H&E stained. The H&E stained KPC and KPCS samples were analyzed for disease grading by two separate pathologists to indicate the stage of the disease. The sample was noted to have normal, PanIN 1, PanIN 2, PanIN 3, or PDA in the pancreas tissue. KPC mice sample sizes were n= 4, 6, 7, 9, 12, and 6 for weeks 4, 8, 12, 16, 24, and 32 respectively. KPCS mice sample sizes were n= 2, 2, 5, 6, 8, and 6 for weeks 4, 8, 12, 16, 24, and 32 respectively. KPC and KPCS PDA samples were examined for the presence of PNI by a pathologist. PNI was defined as previously published^10^.

### Ultrasound Identification and Measurement of Tumor Size

Small animal ultrasound (Vevo770, Visual Sonics) was used to monitor the development and progression of tumors in KPC and KPCS mice starting at 12-16 weeks of age until mouse death. The mice (KPC n=19 and KPCS n=29) were monitored every two weeks to track tumor growth and the day of first tumor detection was determined when the tumor volume exceeded three mm^3^.

### Hemispleen Surgery

The hemispleen surgery was performed as previously described^59^ using wild-type C57/Bl6 mice and tumor cells isolated and grown from KPC and KPCS primary murine tumor samples; time of death for each mouse was recorded.

### Multiplex Immunohistochemistry Analysis

Human PDA specimens were obtained from a published, surgically resected PDA cohort^60^ deposited at the biobank at the Pancreatic Cancer Precision Medicine Center of Excellence at Johns Hopkins according to the Johns Hopkins Medical Institution Institutional Review Board approved protocol IRB0024443. The multiplex immunohistochemistry analysis of this data involved co-registration, visualization and quantitative analysis, as previously described^46,60^. Easily identifiable identical regions (200*200 pixels) from each of the stained image slides were collected, digitized and extracted with Aperio ImageScope software. A customized CellProfiler (Version 2.2.0) pipeline, named “Coregistration 24 markers”, was used to co-register images. Three unique regions of interest (3000*3000 pixels) were extracted from each sample. The regions were selected based on nerve bundle presence within tumor epithelia, confirmed by TUJ1 staining. Two extractions were taken for each region, one including and one directly adjacent to the nerve within the tumor epithelia. The proximal region included tissue within 650 mm of the nerve and the adjacent, distal region included tissue 650-2000 mm from the nerve. No nerve was present in the adjacent region extraction. AEC and color deconvolution functions were run on all images using ImageJ software (64-bit Java 1.8.0_172). This step produced pseudo-colored images that were used for quantification analysis. Images were quantified by a CellProfiler (Version 2.2.0) “Cytometry 17 markers” pipeline. The processed files were then analyzed through FCS Express 6 Image Cytometry software to determine the differences in density of immune cells, defined with CD45 cells, between areas proximal to and distant to a tumor. Gating was performed using stain intensities for M1 TAMs, defined as CD45^+^ CSF1R^+^ CD163^+^, and M2 TAMs, defined as CD45^+^ CSF1R^+^ CD68^+^ CD163^-^. Each gate was established by comparing staining patterns on the dot plot with the original CD45 image. After gating for M1 TAMs and M2 TAMs, populations were measured as a percentage of CD45^+^ cells.

### Quantification and Statistical Analysis

Statistical analysis was performed using GraphPad Prism v9 (Graph Pad Software,La Jolla, CA, USA). The data was presented as means ± SEM. A *P* value of equal or less than 0.05 was considered statistically significant. Statistical significance was calculated between two groups of normally or non-normally distributed data using an unpaired, two-tailed t test or Mann-Whitney U test, respectively. Correlation significance was calculated using a Pearson or Spearman rank correlation test for normally and non-normally distributed data respectively. Statistical analysis of ultrasound tumor size data included log transformation of tumor volume and a linear relationship analysis with a mixed effects model. Western blot densitometry was analyzed though ImageJ software quantification. The mouse survival was analyzed using the Kaplan-Meier method and the log-rank test.

## Key Resources

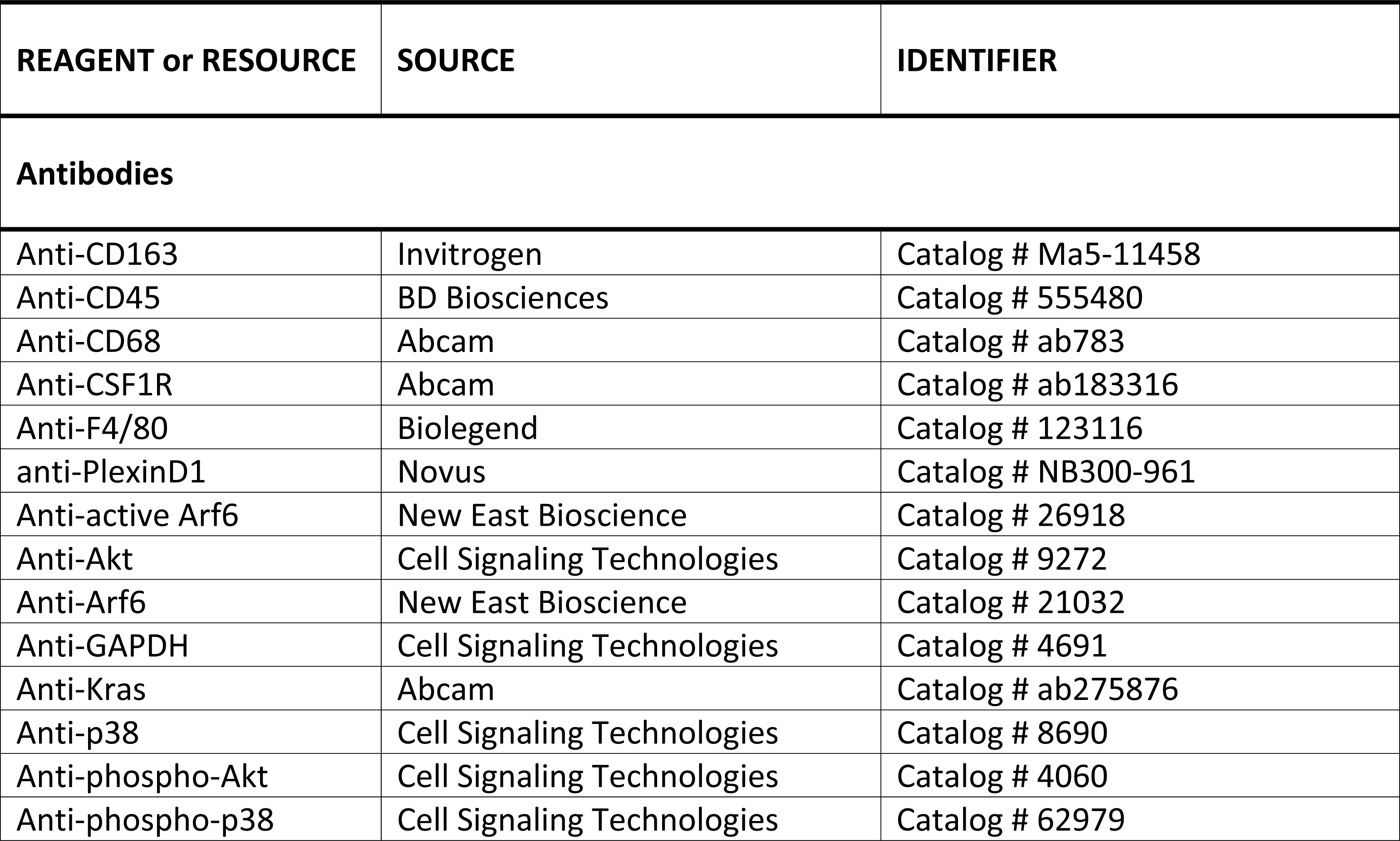
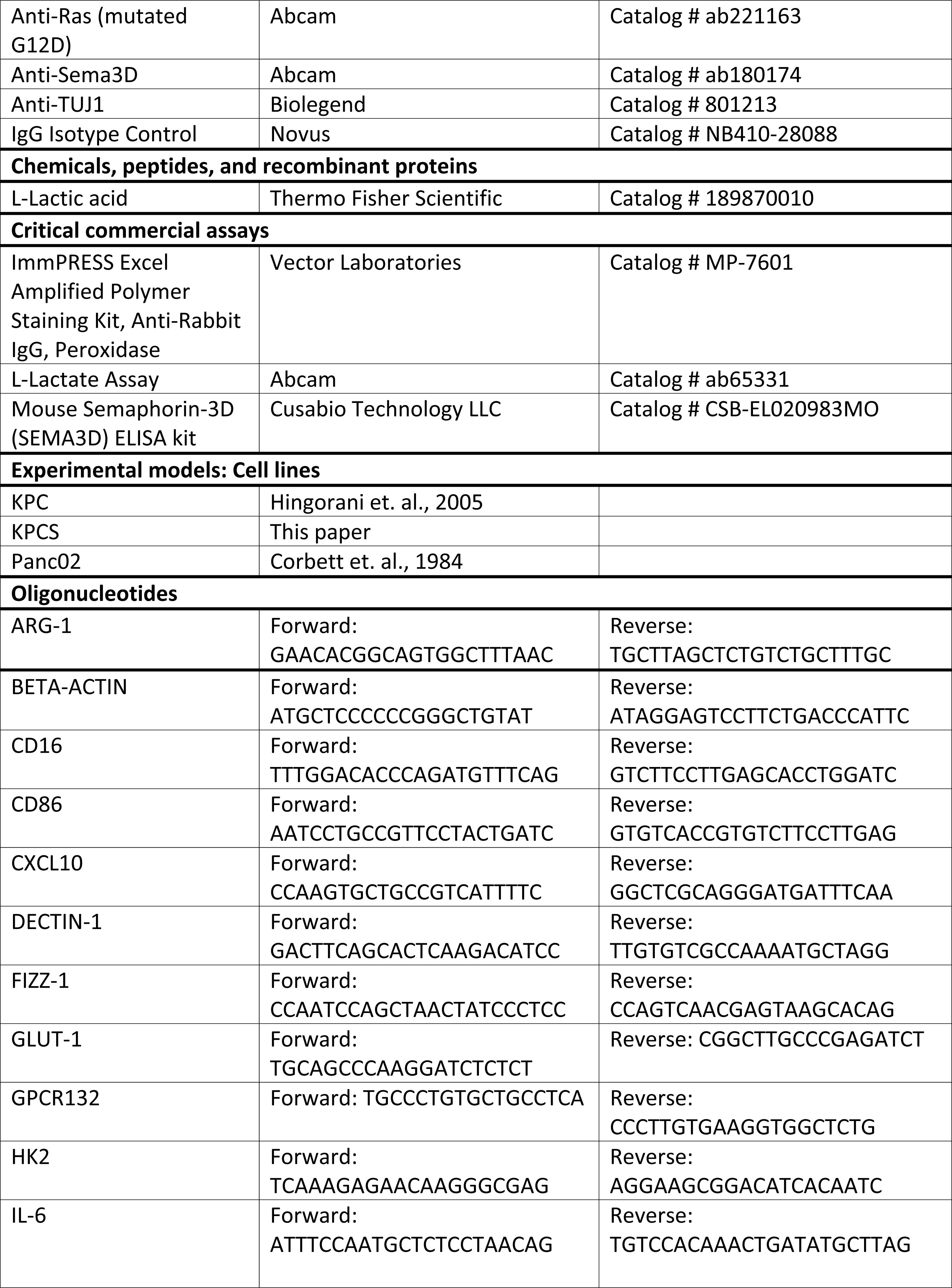
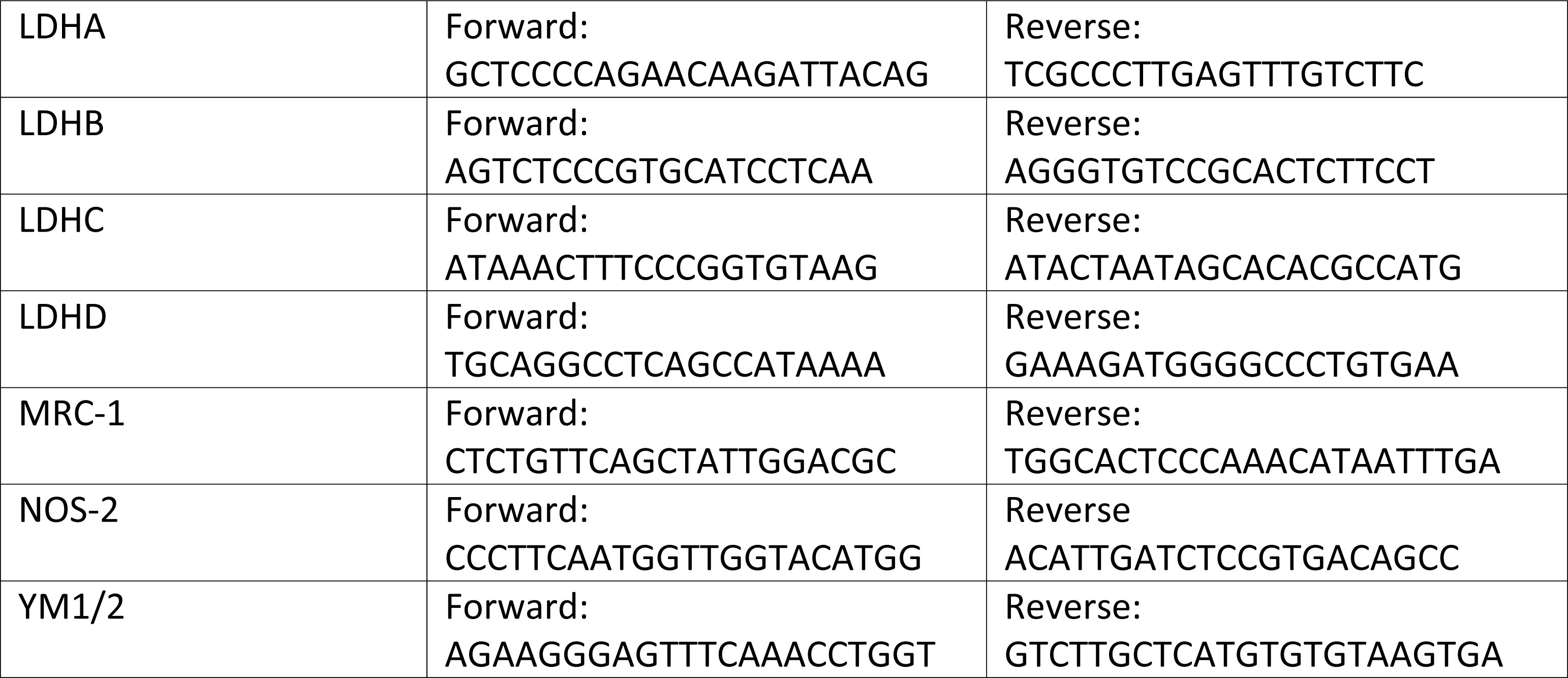

## Data and Software Availability

N/A

## CONTRIBUTIONS

Conceptualization, N.R.J.T., E.M.J., A.K., F.M., L.Z.; Experimentation, N.R.J.T., V.F., S.D., H.E.I., WC.S., J.F., K.L., S.M., X.P., D.T., M.H., K.F., S.S.T. Q.Z., F.M.; Data Analysis, N.R.J.T., S.D. E.T. F.M.; Writing, N.R.J.T., S.D., L.Z.; Supervision, E.M.J., A.K., F.M., L.Z.; Funding, K.F., L.Z.

## ACKNOWLEDGEMENTS

We would like to thank Dr. Yihong Wan for sharing the GPCR132 KO bone marrow for this research. We would also like to thank Dr. Dengfeng Cao at Washington University for providing pathological analysis of our samples. L.Z. was supported by NIH grant R01 CA169702; NIH grant R01 CA197296; and Sidney Kimmel Comprehensive Cancer Center Support Grant P30 CA006973. K.F. was supported as a JSPS Overseas Research Fellow, Japan Society for the Promotion of Science.

## Abbreviations

Pancreatic ductal adenocarcinoma, PDA; Tumor microenvironment, TME; Perineural invasion, PNI; Semaphorin 3D, SEMA3D; Plexin D1, PLXND1; Bone-marrow derived macrophages, BMDMs, Tumor-associated macrophages, TAMs, Pancreatic intraepithelial neoplasia, PanIN; G- protein coupled receptor 132, GPCR132

