## Supplemental Figures for "Tumor- and Nerve-Derived Axon Guidance Molecule Promotes Pancreatic Ductal Adenocarcinoma Progression and Metastasis through Macrophage Reprogramming"

*Supplemental Figure 1:* ***KPC and KPCS mice demonstrate similar rate of tumor growth, related to Figure 1***

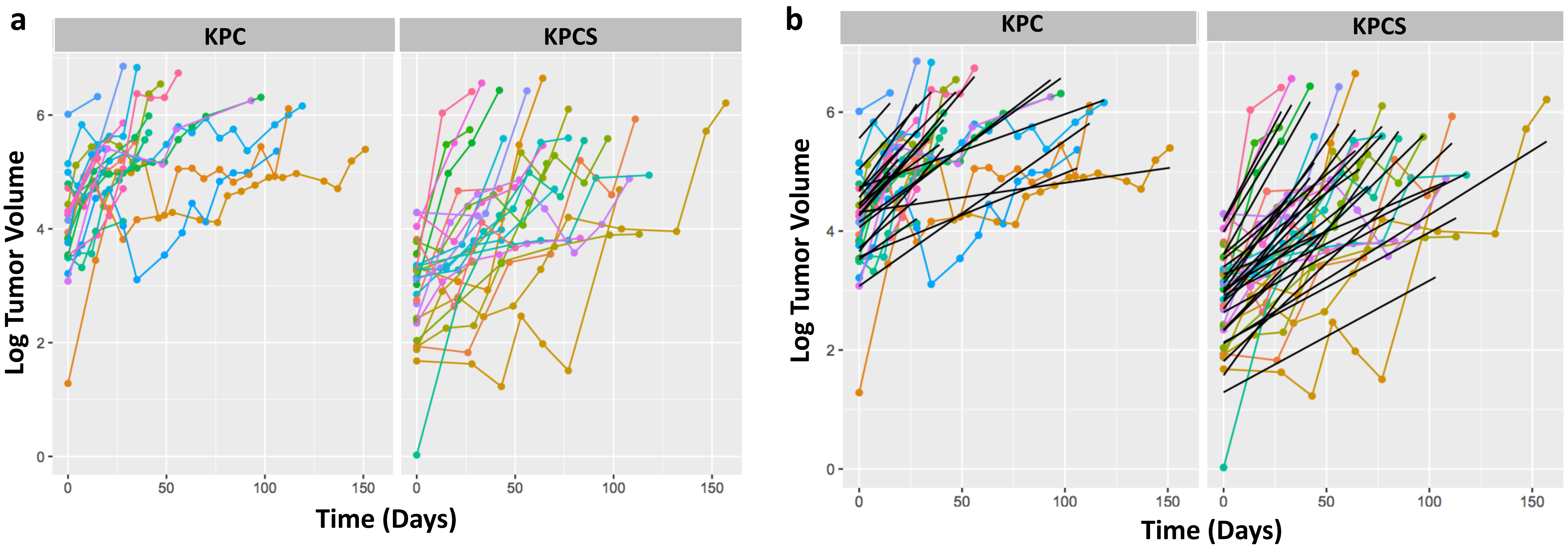

**Supplemental Figure S1:** ***KPC and KPCS mice demonstrate similar rate of tumor growth, related to Figure 1***

a. Spaghetti plots of log-transformed tumor volumes measured over time in KPC and KPC mice.  b. Tumor growth curves for each individual KPC and KPCS mouse based on log-transformed tumor volumes measured over time.

*Supplemental Figure 2:* ***KPC and KPCS mice demonstrate similar nerve density, related to Figure 2***

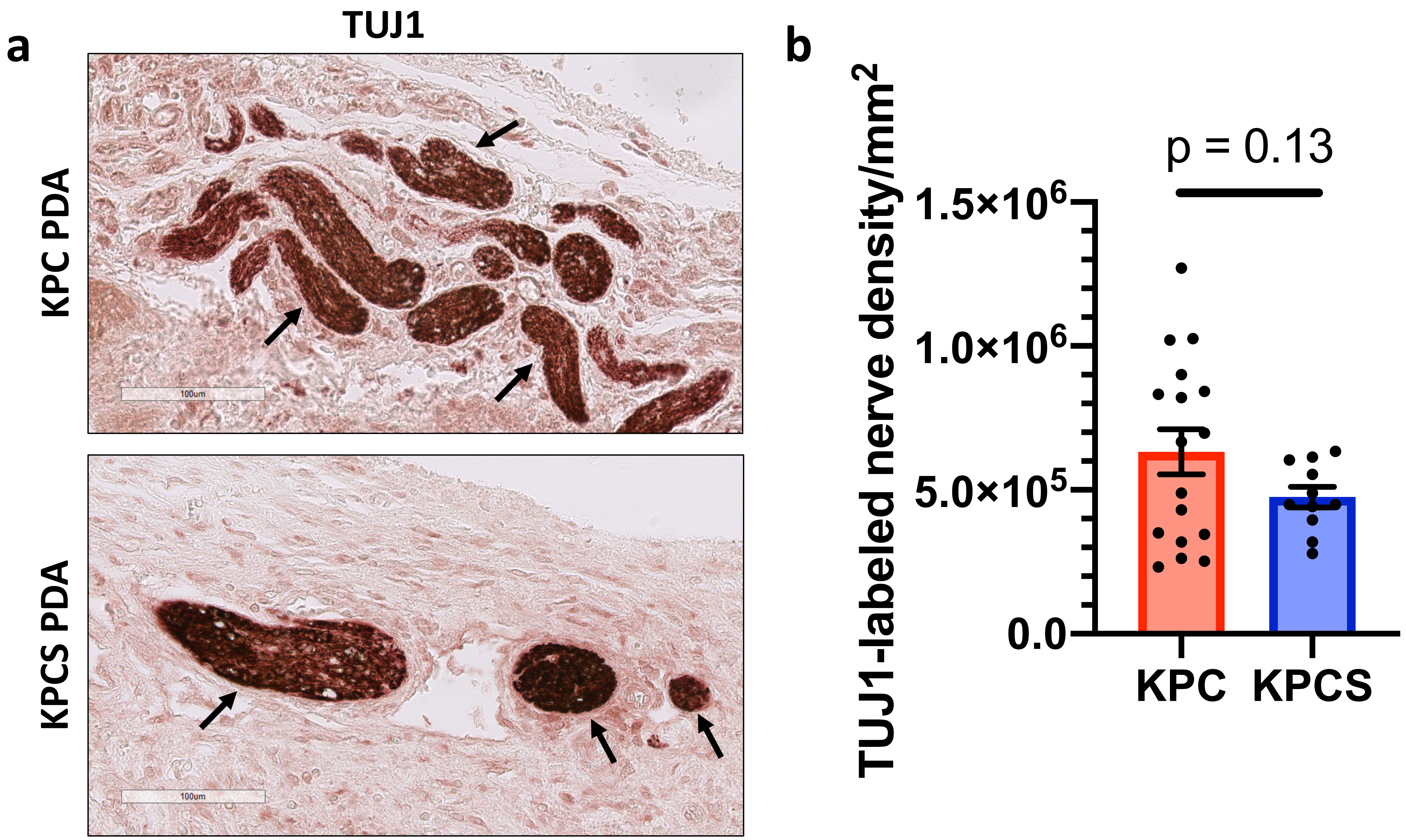

**Supplemental Figure S2:** ***KPC and KPCS mice demonstrate similar nerve density, related to Figure 2***

a. Representative immunohistochemical staining of nerves with anti-TUJ1 antibodies in KPC and KPCS PDAs. Arrows indicate TUJ1-labeled nerve bundles. b. Numbers of TUJ1 labeled nerves per $mm^2$ in the tumor microenvironment of KPC and KPCS mice.

*Supplemental Figure 3:* ***SEMA3D does not reprogram macrophages directly, related to Figure 3***

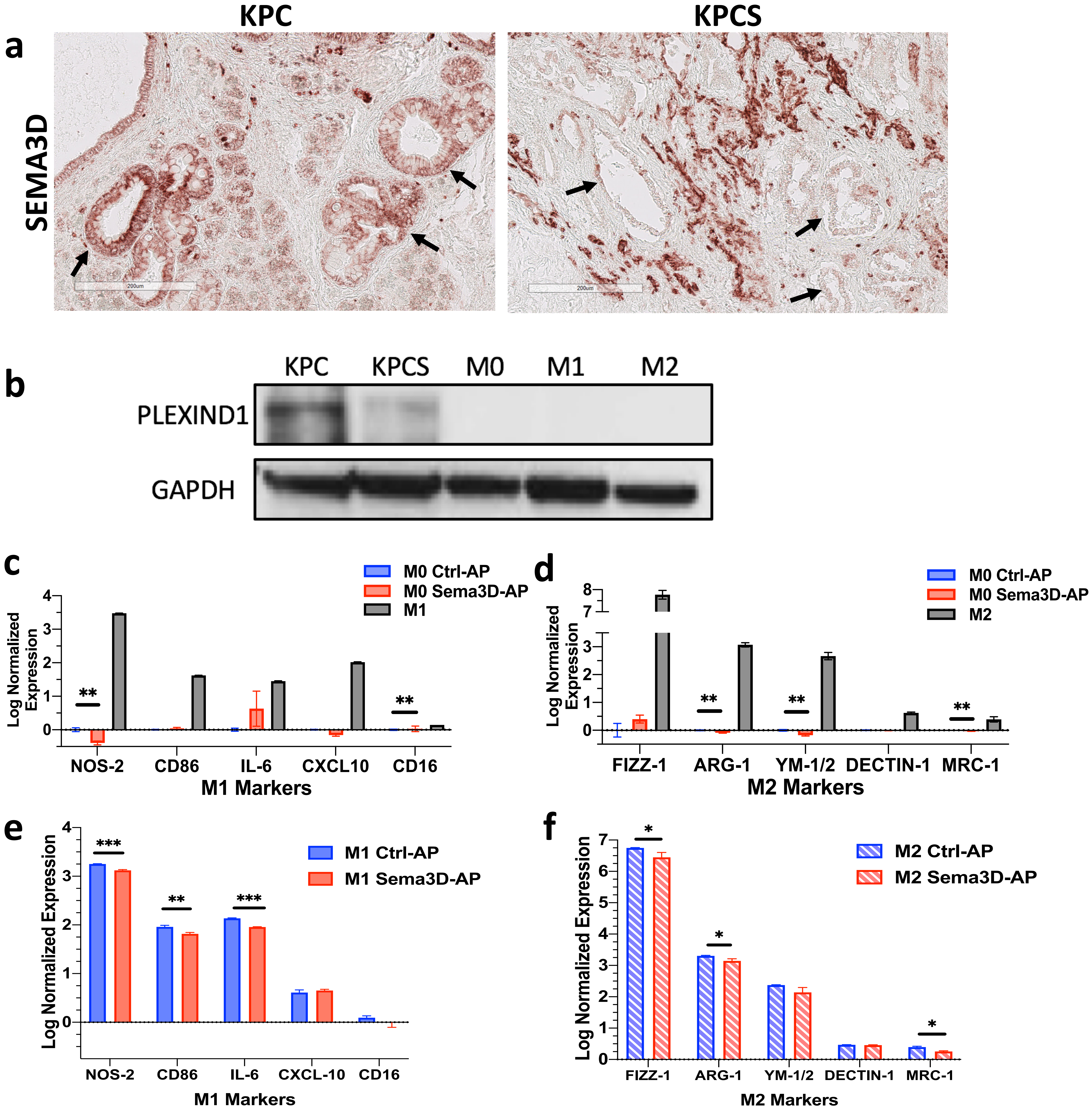

**Supplemental Figure 3:** ***SEMA3D does not reprogram macrophages directly, related to Figure 3***

a. Representative images of immunohistochemical staining of SEMA3D in KPC and KPCS PDA, respectively. Arrows point to PDA tumor cells. b. PLXND1 protein expression in KPC tumor cells (KPC), KPCS tumor cells (KPCS), M0, M1, and M2 macrophages, respectively. c,d. Non-polarized M0 macrophages were treated with CTRL-AP or SEMA3D-AP recombinant protein and subjected to the analysis of gene expression markers for M1 (c) and M2 (d) polarization. Gene expression of polarized M1 and M2 macrophages are shown for comparison. e,f. Polarized M1 (e) and M2 (f) macrophages were treated with CTRL-AP or SEMA3D-AP recombinant protein, respectively, and subjected to the analysis of gene expression markers for M1 and M2 polarization.

*Supplemental Figure 4: Goes with figure 3:* ***SEMA3D reprograms macrophages indirectly through ARF6 signaling and lactate production in PDA tumor cells, related to figure 3***

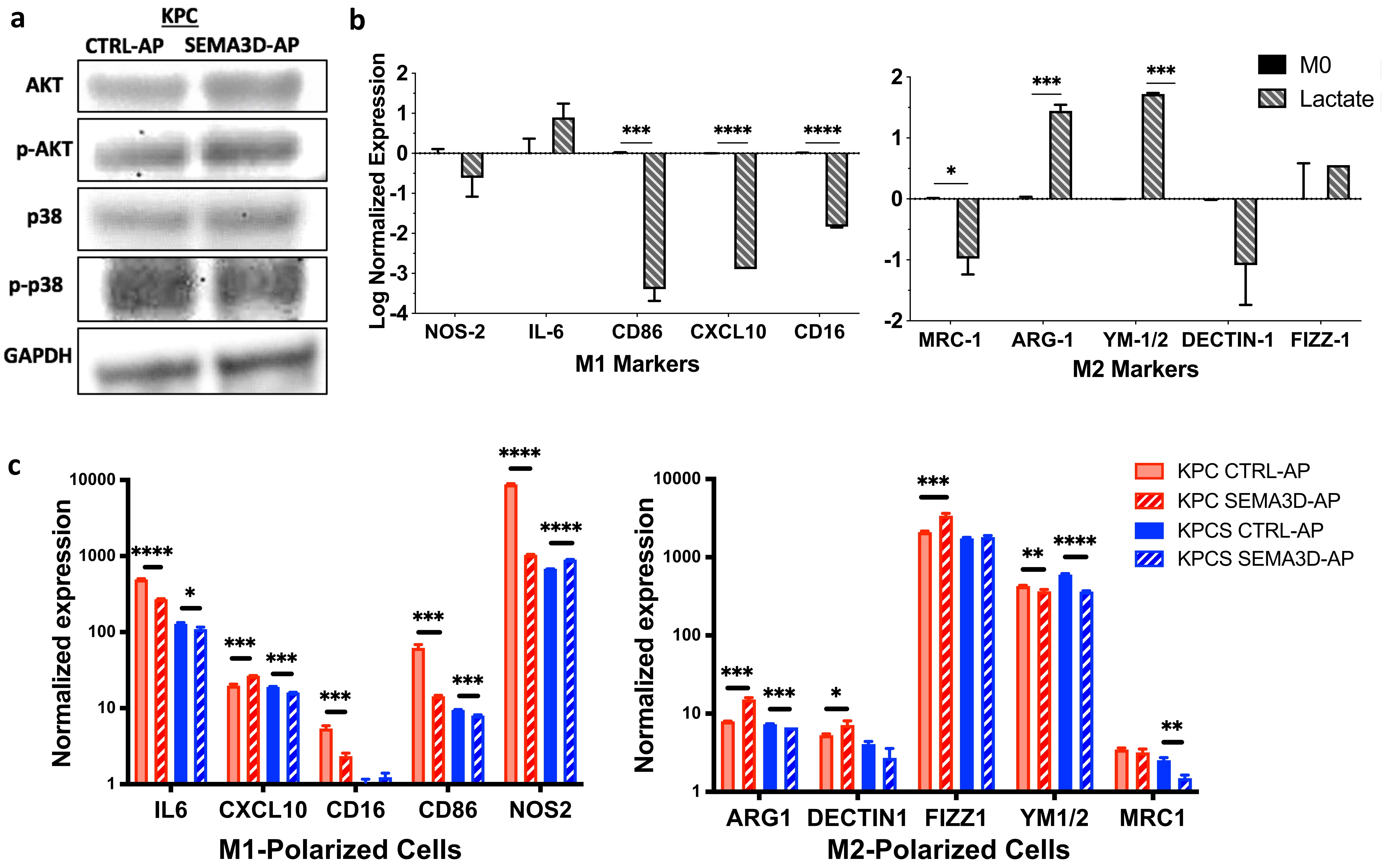

**Supplemental Figure S4:** ***SEMA3D reprograms macrophages indirectly through ARF6 signaling and lactate production in PDA tumor cells,*** **related to Figure 3**
a. Western blot analysis of the expression of AKT, phosphorylated-AKT, p38, phosphorylated p38, and GAPDH proteins in KPC cells treated with CTRL-AP and SEMA3D-AP recombinant protein, respectively. b. Log normalized qRT-PCR measurements of gene expression markers for M1 and M2 polarization in non-polarized M0 macrophages treated with control or 25nM lactate-containing media.
c. qRT-PCR analysis of gene expression markers for M1 and M2 polarization in polarized M1 or M2 macrophages after co-culture with KPC or KPCS cells that have been treated with CTRL-AP or SEMA3D-AP recombinant protein, respectively.

### *Supplemental Figure 5:* ***SEMA3D-induced lactate is sensed by macrophage expressed GPCR132, related to figure 3***

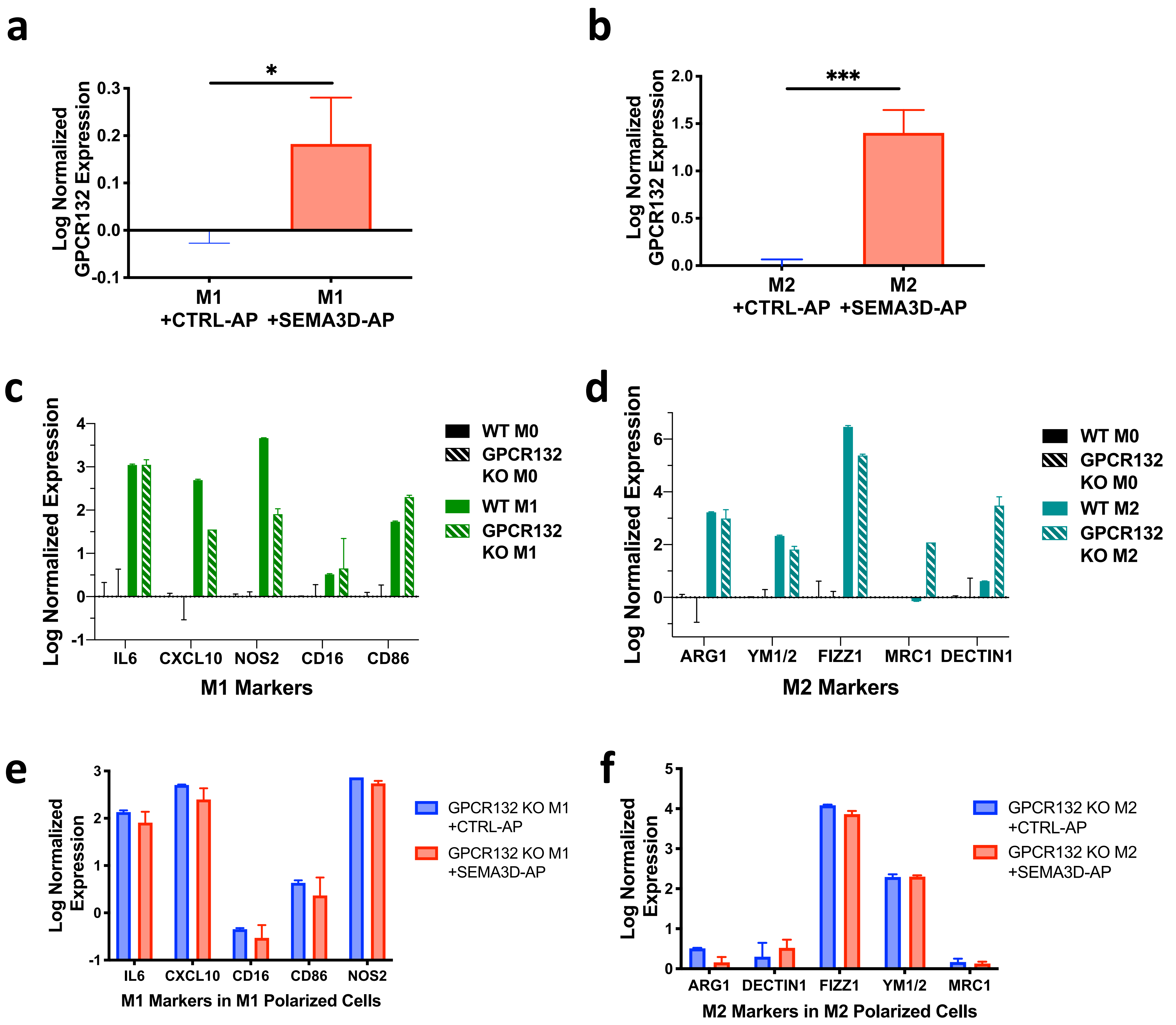

**Supplemental Figure S5:** ***SEMA3D-induced lactate is sensed by macrophage expressed GPCR132, related to Figure 3***

a,b. Log-normalized qRT-PCR measurements of the gene expression of GPCR132 in M1-(a) and M2-(b) polarized macrophages co-cultured with KPC cells that have been treated with CTRL-AP or SEMA3D-AP recombinant protein, respectively. c. qRT-PCR analysis of gene expression markers for M1 polarization in M0 or M1-polarized BMDMs derived from WT or GPCR132 KO mice, respectively. d. qRT-PCR analysis of gene expression markers for M2 polarization in M0 or M2-polarized BMDMs derived from WT or GPCR132 KO mice, respectively. e,f. qRT-PCR analysis of gene expression markers for M1 (e) and M2 (f) polarization in M1- or M2- polarized cells, respectively, co-cultured with KPC cells treated with CTRL-AP or SEMA3D-AP recombinant protein.
